# KMT2D regulates tooth enamel development

**DOI:** 10.1101/2024.08.20.608898

**Authors:** Jung-Mi Lee, Hunmin Jung, Qinghuang Tang, Liwen Li, Soo-Kyung Lee, Jae W. Lee, Yungki Park, Hyuk-Jae Edward Kwon

## Abstract

Amelogenesis, the process of enamel formation, is tightly regulated and essential for producing the tooth enamel that protects teeth from decay and wear. Disruptions in amelogenesis can result in amelogenesis imperfecta, a group of genetic conditions characterized by defective enamel, including enamel hypoplasia, marked by thin or underdeveloped enamel. Mutations in the *KMT2D* (*MLL4*) gene, which encodes a histone H3-lysine 4-methyltransferase, are associated with Kabuki syndrome, a developmental disorder that can involve dental anomalies such as enamel hypoplasia. However, the specific role of KMT2D in amelogenesis remains poorly understood. To address this gap, we generated a conditional knockout mouse model with ectoderm-specific deletion of *Kmt2d* (*Krt14-Cre;Kmt2d^fl/fl^*, or *Kmt2d*-cKO) and characterized the resulting enamel defects using gross, radiographic, histological, cellular, and molecular analyses. Micro-computed tomography and scanning electron microscopy revealed that adult *Kmt2d*-cKO mice exhibited 100% penetrant amelogenesis imperfecta, characterized by hypoplastic and hypomineralized enamel, partially phenocopying human Kabuki syndrome. Additionally, *Kmt2d*-cKO neonates developed molar tooth germs with subtle cusp shape alterations and mild delays in ameloblast differentiation at birth. RNA-seq analysis of the first molar tooth germ at birth revealed that 33.7% of known amelogenesis-related genes were significantly downregulated in the *Kmt2d*-cKO teeth. Integration with KMT2D CUT&RUN-seq results identified 8 overlapping genes directly targeted by KMT2D. Re-analysis of a single-cell RNA-seq dataset in the developing mouse incisors revealed distinct roles for these genes in KMT2D-regulated differentiation across various cell subtypes within the dental epithelium. Among these genes, *Satb1* and *Sp6* are likely direct targets involved in the differentiation of pre-ameloblasts into ameloblasts. Taken together, we propose that KMT2D plays a crucial role in amelogenesis by directly activating key genes involved in ameloblast differentiation, offering insights into the molecular basis of enamel development and related dental pathologies.

## Introduction

Enamel formation, or amelogenesis, is a complex and highly regulated process involving the differentiation and function of ameloblasts, specialized cells responsible for producing the enamel matrix. This process is orchestrated by the enamel organ, the epithelial component of the developing tooth germ. The enamel organ initially undergoes morphogenesis through the bud and cap stages, eventually reaching the bell stage, at which it differentiates into four major cell subtypes: the outer enamel epithelium (OEE), stellate reticulum (SR), stratum intermedium (SI), and inner enamel epithelium (IEE). At the end of the bell stage, the IEE cells further differentiate into ameloblasts, marking the onset of amelogenesis. Amelogenesis is divided into three stages based on ameloblast differentiation and function: presecretory, secretory, and maturation stages, each meticulously regulated by a network of genetic and epigenetic factors (Hu et al. 2007; Lacruz et al. 2017). Disruptions in this process can lead to amelogenesis imperfecta, a condition characterized by hypoplastic and/or hypomineralized enamel, resulting in heightened susceptibility to caries, injury, hypersensitivity, and aesthetic concerns, ultimately impacting oral health. Understanding the regulation of amelogenesis is therefore crucial for maintaining dental health and preventing enamel-related defects.

Recent research underscores the crucial role of epigenetic regulation in orofacial organogenesis, particularly the significance of histone modifications in controlling gene expression essential for tooth development (Deng et al. 2020; Kobayashi et al. 2021; Li et al. 2021; Takagiwa et al. 2024; Yu et al. 2022; Zhang et al. 2023; Zhu et al. 2024). Among these, histone-lysine *N*-methyltransferase 2D (KMT2D), also known as Mixed lineage leukemia 4 (MLL4), catalyzes the methylation of histone H3 lysine 4 (H3K4), a modification associated with active gene transcription (Froimchuk et al. 2017). Mutations in *KMT2D* are associated with Kabuki syndrome (KS), a developmental disorder characterized by distinctive facial features, dental anomalies—including hypodontia, enamel hypoplasia, and delayed tooth eruption— and other systemic defects (Boniel et al. 2021; Matsune et al. 2001; Porntaveetus et al. 2018). However, despite evidence implicating KMT2D in organogenesis, its specific role in amelogenesis and the mechanisms underlying the associated dental anomalies remain poorly understood. A recent study using RNA interference-based knockdown in LS8 cells, an ameloblast-like mouse cell line, have suggested that KMT2D promotes cell proliferation and self-renewal activity via Wnt/β-Catenin signaling (Pang et al. 2021). However, no *in vivo* studies have comprehensively explored the role of KMT2D in tooth development, likely due to the early embryonic lethality of germline *Kmt2d^−/−^* mice, which die around embryonic day (E) 9.5, well before tooth development begins (Lee et al. 2013).

In this study, we define a novel role of KMT2D in amelogenesis by characterizing tooth development in an ectoderm-specific *Kmt2d* conditional knockout (cKO) mouse model. Using gross, radiographic, and microscopic analyses, as well as RNA-sequencing (RNA-seq) and CUT&RUN-sequencing (CUT&RUN-seq), we identified critical genes directly targeted by KMT2D during ameloblast differentiation. These findings provide essential insights into the epigenetic regulation of amelogenesis, highlighting the pivotal role of KMT2D in normal enamel formation and offering potential therapeutic avenues for amelogenesis-related disorders.

## Materials and Methods

### Mouse Strains

*Krt14-Cre* (originally described as ‘*K14*-Cre’) (Andl et al. 2004), and *Kmt2d^fl/fl^* (Lee et al. 2013) mouse lines were used for generating an ectoderm-specific *Kmt2d* cKO model. Mice were maintained by intercrossing or by crossing with C57BL/6J inbred mice. All animal procedures were conducted in accordance with the the Institutional Animal Care and Use Committee at the University at Buffalo (protocol number: ORB16117N). This study is compliant with the ARRIVE (Animal Research Reporting of In Vivo Experiments) guidelines. Additional details on experimental procedures can be found in the Appendix (Lee et al. 2022).

## Results

### KMT2D Deficiency in the Dental Epithelium Causes Amelogenesis Imperfecta

To investigate the role of KMT2D in tooth enamel development, we generated a dental epithelium-specific *Kmt2d*-cKO mouse model (*Kmt2d^fl/fl^;Krt14-Cre*). These mice were born without visible craniofacial anomalies— the palate was fused, and the maxillary and mandibular jaws formed normally (Appendix Fig. 1A). Despite exhibiting scaly skin with scattered hair thinning and reduced body weights compared to controls (*Kmt2d^fll+^*or *Kmt2d^fl/fl^*), *Kmt2d*-cKO mice were viable and appeared otherwise healthy (Appendix Fig. 1B–D) (Egolf et al. 2021). Upon examining the teeth in adult mice (at 8 weeks), we observed that *Kmt2d*-cKO mice displayed chalky-white incisors with rough surfaces, in contrast to the smooth, yellow incisors in control mice (Fig. 1A, E), indicating defective enamel formation. Micro-computed tomography (CT) confirmed significantly reduced enamel mineral density and thickness in both incisors and molars of *Kmt2d*-cKO mice (Fig. 1B, C, F, G, Q, Appendix Fig. 2). Scanning electron microscopy (SEM) of molars further revealed blunt cusp tips, irregular enamel surfaces, and poorly formed enamel rods in the *Kmt2d*-cKO mice (Fig. 1D, H, R). To determine whether reduced enamel thickness in the *Kmt2d*-cKO molars, particularly in the cusp tips, was due to defective formation or post-eruption wear, we analyzed pre-eruptive molars at 2 weeks. Consistent with the findings at 8 weeks, pre-eruptive molars in *Kmt2d*-cKO mice at 2 weeks exhibited significantly reduced enamel mineral density and thickness with hypoplastic tooth crowns; however, molar cusp tips were not blunt at this stage (Fig. 1I–Q). Collectively, these findings demonstrate that *Kmt2d* deficiency leads to a combination of hypoplastic and hypomineralized amelogenesis imperfecta, highlighting its essential role in ensuring proper enamel thickness and mineral density during amelogenesis. Moreover, this study suggests that the *Kmt2d*-cKO strain represents a valuable model for studying amelogenesis and the etiology of dental anomalies associated with KS.

**Figure 1.**
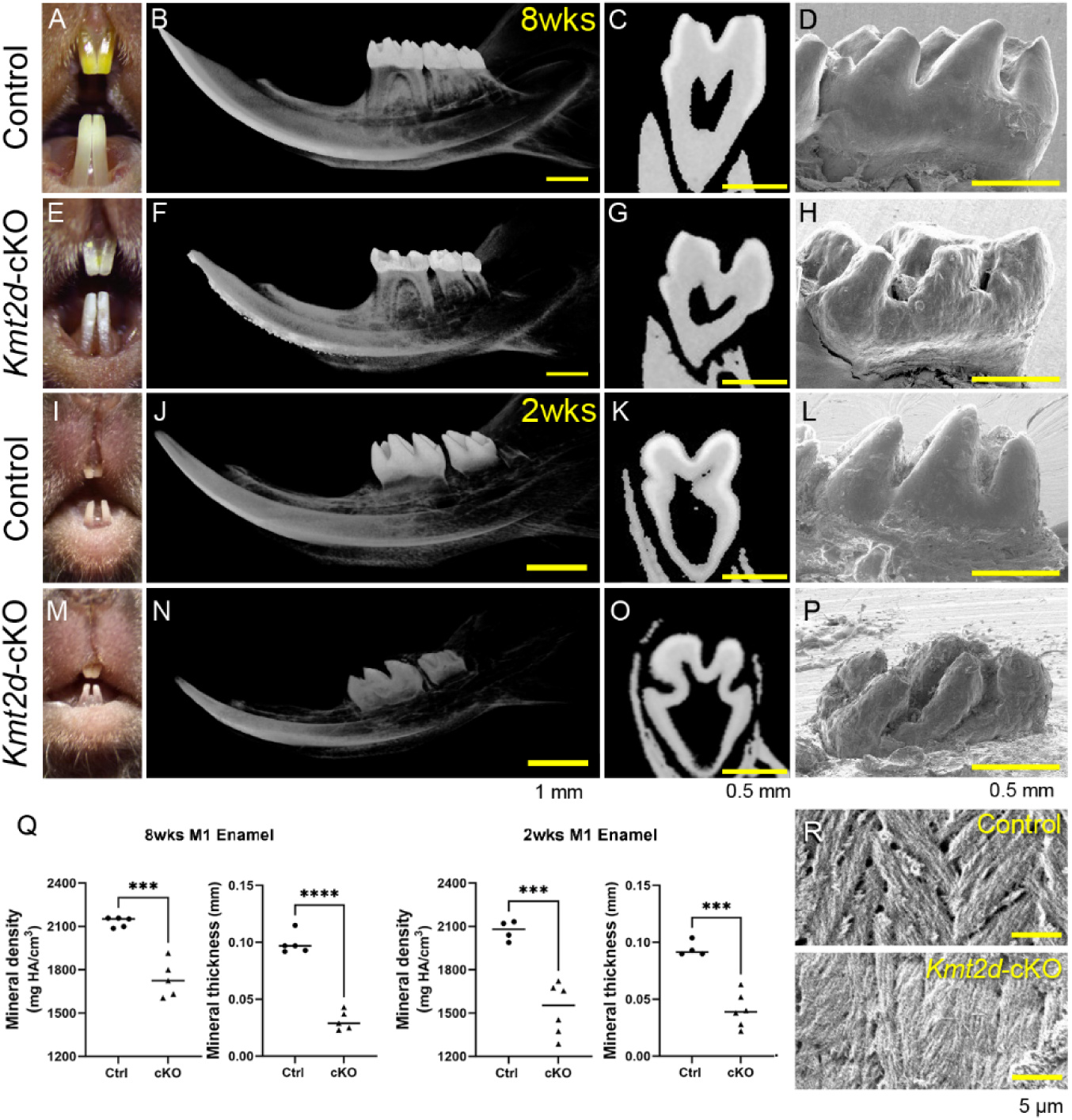
*Kmt2d*-cKO mice exhibit defective enamel, phenocopying human Kabuki syndrome. (A–D, E– H, I–L, M–P) Comparison of control and dental epithelium-specific conditional *Kmt2d*-deficiency (*Kmt2d*-cKO) mouse teeth at 8 weeks (A–H) and 2 weeks (I–P). *Kmt2d*-cKO mice exhibited chalky-white incisors with a rough surface (E) compared to control incisors, which were yellow and had a smooth surface (A), at 8 weeks. Micro-CT analysis of the mandible showed a defective enamel layer with reduced radiopacity and thickness in the *Kmt2d*-cKO incisor and molar teeth at both 8 and 2 weeks (F, G, N, O), compared to controls (B, C, J, K). Panels C, G, K, and O are frontal/coronal slices of the first molar in B, F, J, and N. (D, H, L, P) SEM images of the first molar tooth crown from control (D, L) and *Kmt2d*-cKO mice (H, P) showed that *Kmt2d*-cKO teeth exhibit blunt cusp tips at 8 weeks (D, H) and a hypoplastic crown at 2 weeks (L, P). (Q) Quantification of enamel mineral density and thickness in the first molars from control and *Kmt2d*-cKO mice at 8 weeks and 2 weeks. (R) SEM images of enamel rod microstructures in control and *Kmt2d*-cKO mice. Scale bars: 1 mm (in B, F, J, N), 0.5 mm (in C, D, G, H, K, L, O, P), and 5 µm (in R). M1, first molar. \*\*\**p* < 0.001; \*\*\*\**p* < 0.0001. *n* = 4–6 per group.

### KMT2D-cKO Teeth Exhibit Cusp Alterations and Delayed Ameloblast Differentiation During Embryonic Development

To investigate the developmental mechanisms underlying the role of KMT2D in amelogenesis, we examined the histological features of the first molar in control and *Kmt2d*-cKO mice during embryonic tooth development. Hematoxylin and eosin-stained frontal sections of the first molar tooth germ revealed that *Kmt2d*-cKO embryos exhibited nearly normal tooth morphogenesis throughout the cap (E14.5) and bell (E16.5) stages with only minor alterations in cusp shape observed at E16.5 (Fig. 2A–D). At birth (P0.5), when ameloblasts transition from the presecretory to secretory stage, *Kmt2d*-cKO mice displayed slightly shorter but broader cusp tips, measured consistently in sections at the largest dimension of the mesiobuccal ‘protoconid’ and mesiolingual ‘metaconid’ cusps of the first molar (Fig. 2E, F). Higher magnifications revealed that ameloblasts in *Kmt2d*-cKO mice were less polarized and shorter compared to controls (Fig. 2G–J). By P7, corresponding to the late secretory stage, *Kmt2d*-cKO mice displayed reduced enamel matrix protein deposition, particularly along the lateral molar surfaces (Fig. 2K, L, black arrow), while cusp regions remained comparable to controls. This finding suggests a delay in functional ameloblast differentiation due to *Kmt2d* deletion. By P14, during the late maturation stage, *Kmt2d*-cKO mice exhibited a generalized reduction in enamel thickness in the first molar (Fig. 2M, N). These results indicate that KMT2D is critical for timely ameloblast differentiation, which supports proper enamel formation. To explore the cellular mechanisms underlying these developmental phenotypes, we analyzed cell proliferation using EdU staining and apoptosis using TUNEL staining at E18.5. *Kmt2d*-cKO mice exhibited increased cell proliferation in both the pre-ameloblast and pre-odontoblast layers, while apoptosis levels remained unchanged (Appendix Fig. 3A–F). This suggests that an overall developmental delay associated with *Kmt2d* deficiency may cause a non-specific increase in cell proliferation within both the dental epithelium and mesenchyme. Further, we assessed KMT2D expression during tooth development using immunofluorescence staining. Consistent expression was detected in the enamel organ (Appendix Fig. 4A–J). In contrast, *Kmt2d*-cKO mice displayed a marked reduction of KMT2D signals in the dental epithelium, validated by real-time qPCR (qRT-PCR) analysis (Appendix Fig. 4K), which confirmed efficient deletion of *Kmt2d* in the epithelium while mesenchymal expression remained unaffected. Interestingly, despite unchanged *Kmt2d* RNA levels in the dental mesenchyme, KMT2D protein signals were reduced in the dental mesenchyme of *Kmt2d*-cKO tooth germs. This observation suggests that epithelial-mesenchymal interactions—known to be critical regulators of dental development—may be disrupted by epithelial *Kmt2d* deficiency, indirectly altering mesenchymal gene expression or protein stability through impaired paracrine signaling.

**Figure 2.**
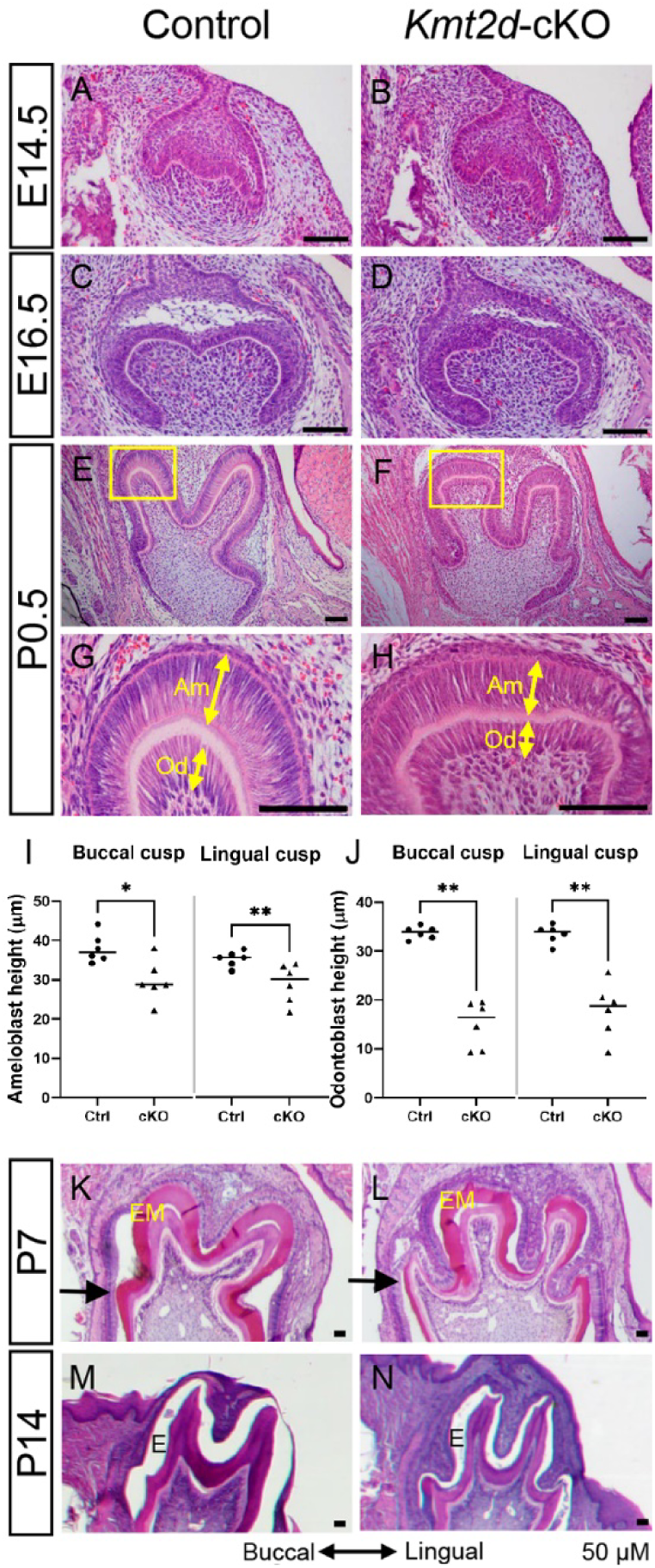
*Kmt2d*-cKO mouse embryos exhibit nearly normal tooth morphogenesis, with minor alterations in cusp shape and ameloblast cell polarization at birth. (A–H, K–N) Hematoxylin and eosin staining of frontal sections of the first molar tooth germ in control (A, C, E, K, M) and *Kmt2d*-cKO mice (B, D, F, L, N) at E14.5 (A, B), E16.5 (C, D), P0.5 (E, F), P7 (K, L), and P14 (M, N). (G, H) Higher magnifications of the buccal cusp (yellow boxes in panels E and F) show broadened cusp tips with shorter ameloblasts and odontoblasts in the *Kmt2d*-cKO mice compared to controls. Scale bars: 50 µm (in panels A–H, K–N). Am, ameloblast; Od, odontoblast; EM, enamel matrix protein; E, enamel. The black double-headed arrow indicates buccal and lingual directions. (I, J) Quantification of ameloblast (I) and odontoblast (J) heights in the buccal and lingual cusps, as shown in panels G and H. \**p* < 0.05; \*\**p* < 0.01. *n* = 6 per group.

### KMT2D Regulates Amelogenesis-Related Gene Expression During Early Amelogenesis

To elucidate the molecular mechanisms underlying the gross and microscopic changes observed in *Kmt2d*-cKO mouse teeth, we performed genome-wide mRNA expression analysis (RNA-seq) using the first molars from control and *Kmt2d*-cKO mice at birth (P0.5). Our analysis identified 2,366 differentially expressed genes (DEGs), including 1,485 downregulated and 881 upregulated genes, in the *Kmt2d*-cKO group compared to controls (adjusted *p*-value ≤ 0.001, log_2_FC ≥ 0.4) (Fig. 3A). Over-representation analysis (ORA) and gene set enrichment (GSE) analysis of these DEGs revealed significant enrichment of gene ontology (GO) terms related to “biomineral tissue development,” “odontogenesis,” “tooth mineralization,” and “amelogenesis” (Fig. 3B; Appendix Fig. 5). Interestingly, “positive regulation of cell cycle process” was also significantly enriched among the upregulated genes, aligning with the observed increase in cell proliferation in *Kmt2d*-cKO mice (Appendix Fig. 5). Given KMT2D’s established role in transcriptional activation, we focused on the downregulated DEGs to identify genes relevant to amelogenesis. By intersecting the 1,485 downregulated DEGs with a curated list of 92 amelogenesis-related genes (hereafter “amelogenesis genes”) from previous studies (Appendix Table 1) (Bloch-Zupan et al. 2023; Dong et al. 2023; Smith et al. 2017; Wright et al. 2015), we found that 33.7% (31 of 92) of the amelogenesis genes were significantly downregulated in *Kmt2d*-cKO mice (Fig. 3C). These genes included key enamel matrix protein genes, such as amelogenin (*Amelx*), ameloblastin (*Ambn*), and enamelin (*Enam*) (Fig. 3D). The downregulation of these genes was further validated by qRT-PCR (Fig. 3E; Appendix Fig. 6A). Collectively, these results demonstrate that KMT2D is critical for early amelogenesis, potentially through the timely activation of gene regulatory programs essential for proper ameloblast differentiation.

**Figure 3.**
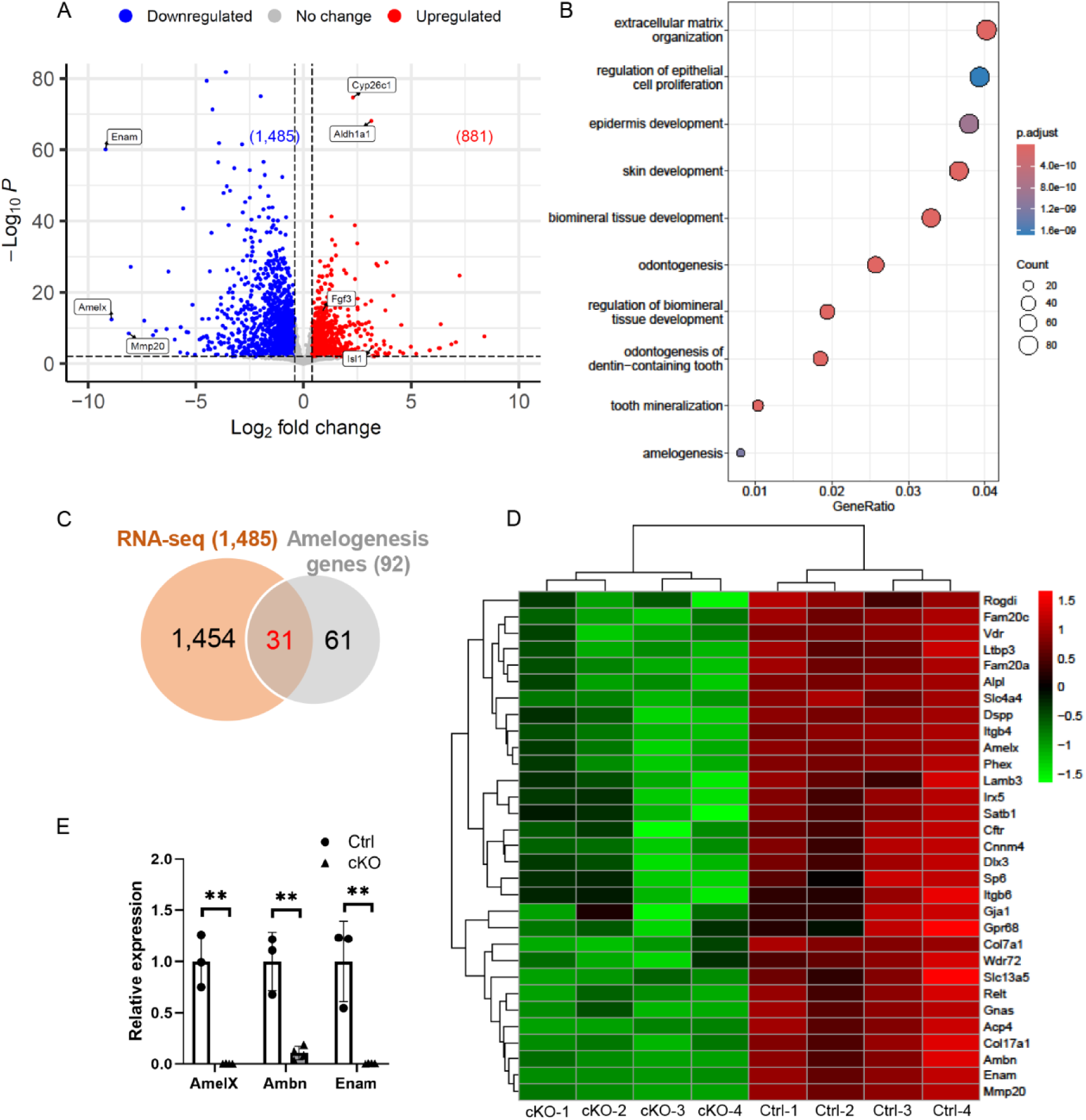
KMT2D regulates amelogenesis-related gene expression in the molar tooth germ. (A–D) Genome-wide mRNA expression analysis in control and *Kmt2d*-cKO first molars at P0.5. (A) Volcano plot showing a total of 2,366 differentially expressed genes (DEGs)—881 upregulated and 1,485 downregulated (adjusted *p*-value ≤ 0.001, log_2_FC ≥ 0.4). (B) Enriched gene ontology (GO) biological processes from over-representation analysis (ORA) of the significant DEGs. (C) Integration of the downregulated DEGs and 92 genes related to amelogenesis, showing that 33.7% (31 of 92) of amelogenesis genes overlap with the downregulated DEGs. (D) Heatmap of the 31 overlapping genes identified in panel C. (E) Real-time qPCR results for key amelogenesis genes (*Amelx, Ambn,* and *Enam*) selected from these 31 overlapping genes, validating the significantly decreased expression in the *Kmt2d*-cKO first molar compared to control. \*\**p* < 0.01. *n* = 3–4 per group.

### Identification of 8 Candidate Amelogenesis Genes Directly Targeted by KMT2D

Next, to identify the direct transcriptional target genes of KMT2D, we mapped genome-wide KMT2D-binding loci using KMT2D CUT&RUN-seq on wildtype P0.5 mandibular first molars. Across two replicates (replicate #1: 4,198 peaks; replicate #2: 3,030 peaks), we identified 2,271 overlapping peaks, which were used to minimize false-positive results. Approximately 50% of these peaks were located in promoter regions, while about 47% were found in potential enhancer regions, including intronic, intergenic, and immediate downstream regions (Fig. 4A). Using the GREAT tool for GO enrichment analysis, we found that “histone H3 acetylation,” “histone methylation,” and “enamel mineralization” were among the most enriched biological processes, suggesting that KMT2D may regulate amelogenesis through its role as a histone H3K4 methyltransferase (Fig. 4B). To further link KMT2D-binding loci to amelogenesis, we integrated the CUT&RUN-seq data with a list of 92 amelogenesis genes. This analysis revealed that 14% (13 of 92) of the amelogenesis genes were directly bound by KMT2D (Fig. 4C). Finally, by integrating the RNA-seq and CUT&RUN-seq results with the amelogenesis gene list, we identified 8 overlapping genes directly targeted by KMT2D (Fig. 4C, D).

**Figure 4.**
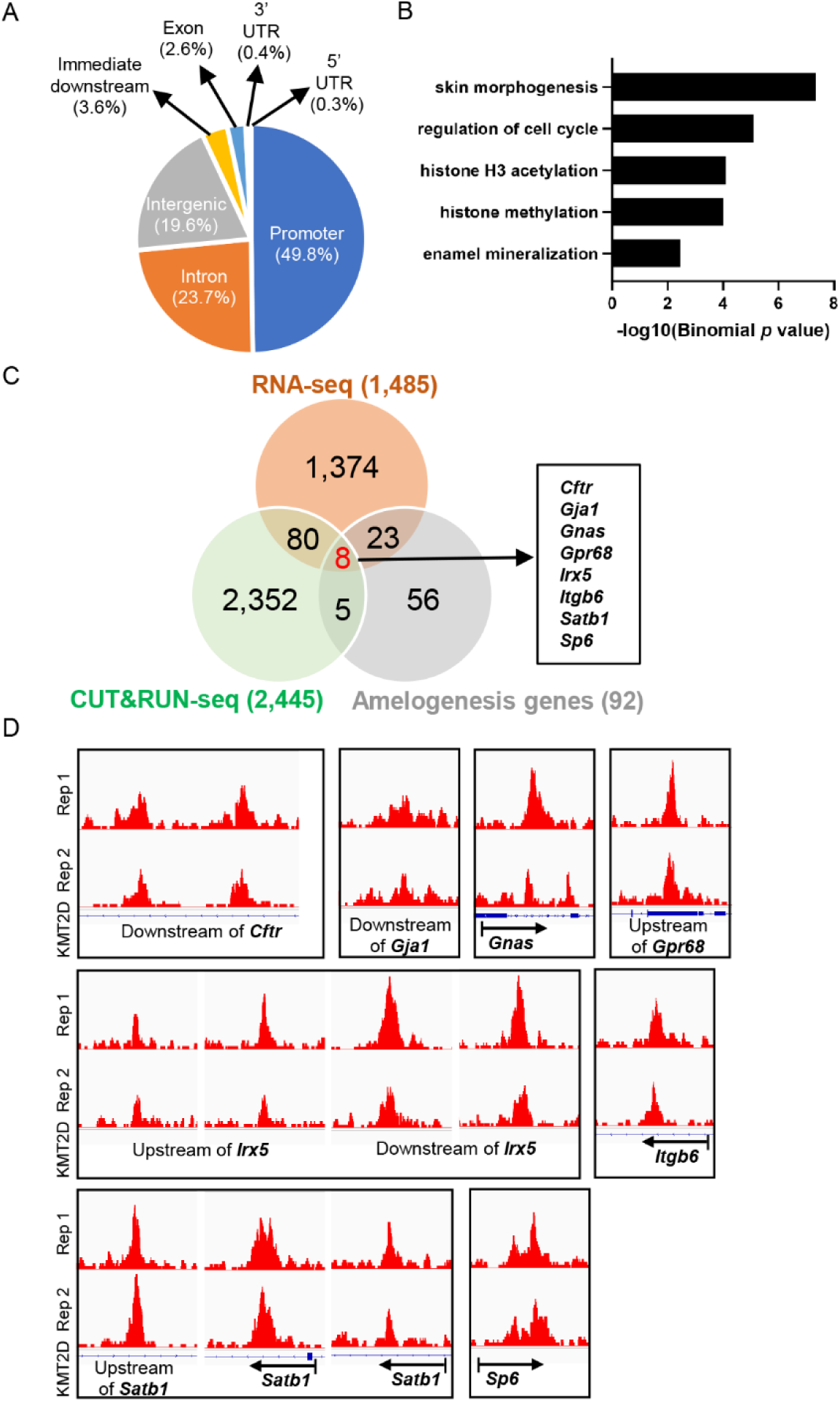
Identification of amelogenesis-related genes directly targeted by KMT2D. (A–D) KMT2D CUT&RUN-seq analysis results in wildtype first molars at P0.5, with two replicates. (A) Distribution pattern of the KMT2D binding peaks across genomic regions. (B) Enriched GO biological processes associated with KMT2D binding peaks using the GREAT tool. (C) Three-way intersection of CUT&RUN-seq-based KMT2D target genes, RNA-seq-based *Kmt2d*-cKO downregulated DEGs (as identified in Figure 3C), and 92 amelogenesis genes, identifying 8 overlapping genes directly targeted by KMT2D. (D) KMT2D binding loci near the identified 8 direct target genes of KMT2D from two CUT&RUN-seq replicates (KMT2D_Rep 1 and KMT2D_Rep 2).

### KMT2D Regulates Distinct Sets of Genes in Different Subtypes of the Enamel Organ During Ameloblast Differentiation

Based on our findings that dental epithelium-specific *Kmt2d* deletion primarily causes delay and defects in ameloblast differentiation (Fig. 2G–J), we compared the mRNA expression patterns of *Kmt2d* and the 8 candidate genes directly targeted by KMT2D—identified through integration of RNA-seq and CUT&RUN-seq results with amelogenesis genes—across the major cell subtypes of the dental epithelial enamel organ. We re-analyzed single-cell RNA-seq datasets from the mouse incisor dental epithelium (Chiba et al. 2020) and classified the enamel organ cells into four major subtypes: inner enamel epithelium/outer enamel epithelium (IEE/OEE), stratum intermedium/stellate reticulum (SI/SR), pre-ameloblast (pre-AmB), and ameloblast (AmB) (Fig. 5A). Pseudotemporal trajectories revealed bifurcation into ameloblast (Fig. 5A, upward arrow) and non-ameloblast lineages (Fig. 5A, downward arrow) from the IEE/OEE, consistent with the current model of amelogenesis (Fresia et al. 2021). *Kmt2d* was expressed ubiquitously across all enamel organ cell subtypes, though not in every individual cell (Fig. 5B). Co-expression analysis of *Kmt2d* and the 8 candidate genes revealed cell type-specific differential expression during dental epithelial cell differentiation in the enamel organ. Notably, *Cftr, Satb1*, and *Sp6* showed strong co-expression with *Kmt2d* in the pre-AmB and AmB subtypes (Fig. 5C, top row), suggesting their potential involvement in KMT2D-regulated pre-AmB-to-Amb differentiation. *Gnas* and *Gpr68* exhibited co-expression with *Kmt2d* across pre-AmB, AmB, and SI/SR subtypes (Fig. 5C, middle row), indicating their potential roles in both KMT2D-regulated AmB and non-AmB differentiation. *Irx5* was predominantly co-expressed with *Kmt2d* in pre-AmB, while *Itgb6* was primarily co-expressed in AmB, and *Gja1* in SI/SR subtypes (Fig. 5C, bottom row). Collectively, these findings indicate that KMT2D regulates distinct sets of genes within specific enamel organ cell subtypes during ameloblast differentiation, underscoring its critical role in the process.

**Figure 5.**
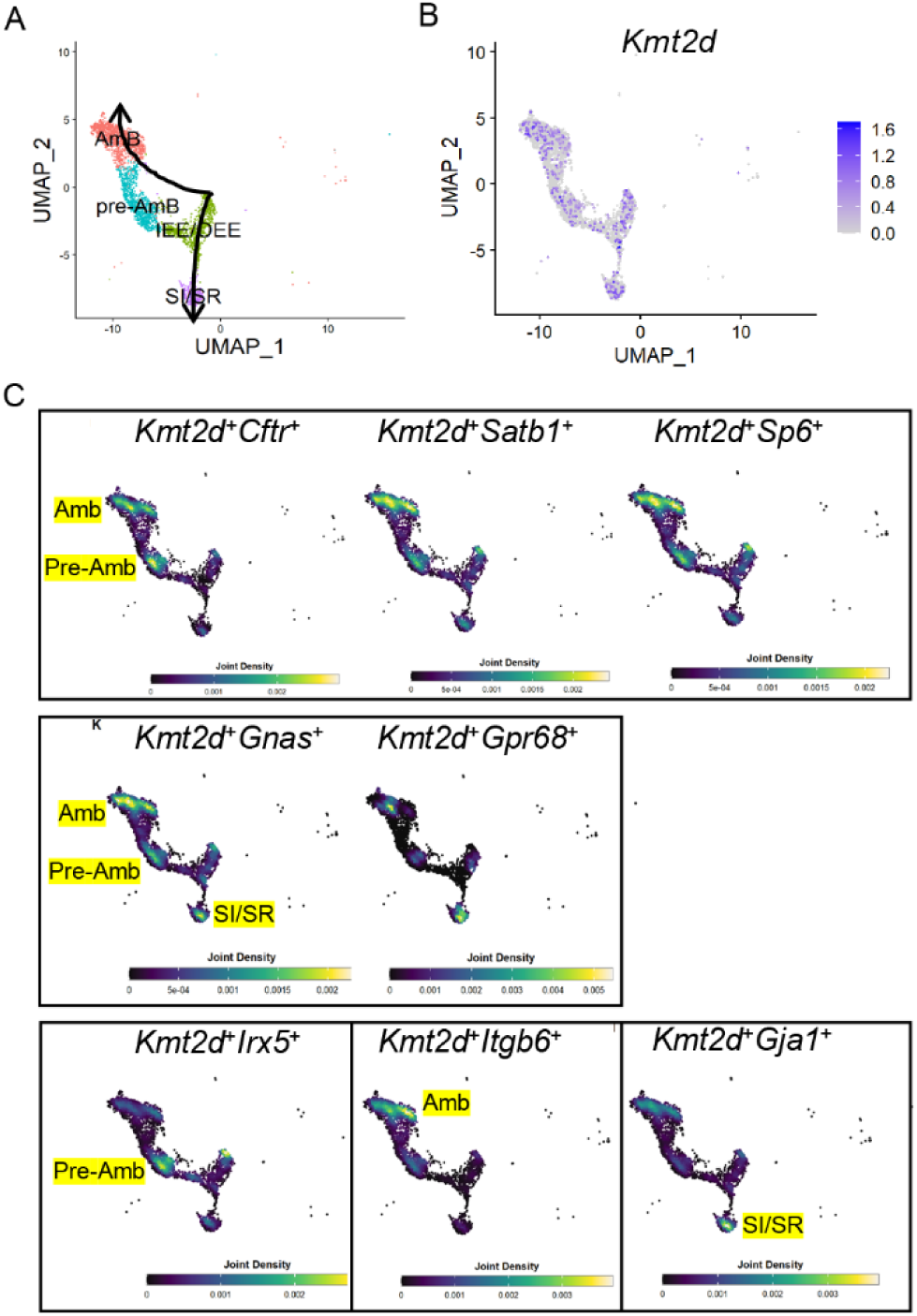
Cell subtype-specific differential gene expression analysis reveals a distinct involvement of KMT2D target genes during the differentiation of dental epithelium. (A–C) Re-analysis of a single-cell RNA-seq dataset from P7 mouse incisors. (A) Uniform Manifold Approximation and Projection (UMAP) showing the classification of the dental epithelial cells into four major cell types: inner enamel epithelium/outer enamel epithelium (IEE/OEE), stratum intermedium/stellate reticulum (SI/SR), pre-ameloblast (pre-AmB), and ameloblast (AmB). The UMAP also indicates the pseudotemporal trajectories of these cell types, with arrows showing the bifurcation of ameloblast and non-ameloblast cells (arrows). (B) Identification of cell subtypes expressing *Kmt2d*. (C) Joint density plots visualizing the co-expression patterns of 8 target genes with *Kmt2d*.

## Discussion

In this study, we established that KMT2D is a crucial factor in amelogenesis, as evidenced by the significant reduction in enamel thickness and mineral density observed in the teeth of *Kmt2d*-cKO mice (Fig. 1), along with defects in ameloblast differentiation (Fig. 2). By integrating CUT&RUN-seq-derived KMT2D target genes (Fig. 4) with RNA-seq-identified downregulated DEGs in *Kmt2d*-cKO mice (Fig. 3) and previously known amelogenesis-related genes, we identified 8 overlapping genes regulated by KMT2D that are implicated in amelogenesis. Further integration of our findings with single-cell transcriptomics datasets revealed the potential distinct roles for these genes in KMT2D-regulated differentiation trajectories within the enamel organ (Fig. 5). We conducted a literature review focusing on mouse and human enamel phenotypes associated with these 8 KMT2D-regulated amelogenesis genes. Given that *Kmt2d*-cKO mice exhibit initial defects in ameloblast differentiation at birth, we primarily focused on the *Cftr*, *Satb1*, and *Sp6* genes. These genes were identified as being involved in the KMT2D-regulated differentiation trajectory from pre-ameloblasts to ameloblasts during early amelogenesis (Fig. 5C, top row). Our findings demonstrate a significant reduction in *Cftr*, *Satb1*, and *Sp6* mRNA expression, as confirmed by qRT-PCR, exclusively in the cKO dental epithelium, with no changes observed in the cKO dental mesenchyme (Appendix Fig. 6A). Additionally, CUT&RUN-qPCR analysis revealed specific fold enrichment of these genes in the dental epithelium compared to the dental mesenchyme (Appendix Fig. 6B), further highlighting the dental epithelium-specific impact of *Kmt2d* deletion.

CFTR, primarily known for its role in cystic fibrosis, functions as a chloride ion channel across the cell membrane. In amelogenesis, *Cftr* is expressed in maturation-stage ameloblasts, with structural defects observed in these regions in *Cftr*-deficient mouse incisors (Bronckers et al. 2010). However, enamel defects in *Cftr*^−/−^ mice are restricted to incisors, with the molars appearing normal (Gawenis et al. 2001). These studies suggest that *Cftr* may not be the primary factor responsible for the early amelogenesis defects observed in our *Kmt2d*-cKO model. Nevertheless, investigating whether CFTR contributes to KMT2D-regulated late-stage amelogenesis, particularly during the maturation stage, remains an intriguing question. Although *Cftr* expression levels were significantly lower compared to *Satb1* and *Sp6* (average RPKM in controls: *Cftr*, 6.56; *Satb1*, 319.30; *Sp6*, 1,076.77), the observed transcriptional changes at P0.5 likely reflect the early impact of *Kmt2d* loss on ameloblasts developmental trajectories, which may subsequently influence the expression of these genes during later amelogenesis stages.

SATB1, a transcription factor and genome organizer, is essential for establishing ameloblast cell polarity and regulating enamel formation (Zhang et al. 2019). *Satb1* expression is highest in presecretory-stage ameloblasts, reduced in secretory-stage ameloblasts, and significantly diminished in maturation-stage ameloblasts. Notably, *Satb1*^−/−^ mice exhibit a tooth phenotype strikingly similar to our *Kmt2d*-cKO mice, with defects in ameloblast differentiation during early amelogenesis. Moreover, Kohlschütter-Tönz syndrome-like (KTZSL) disorder, caused by mutations in *SATB1*, shares developmental defects with KS, including intellectual disabilities, hypotonia, and enamel hypoplasia (den Hoed et al. 2021). However, the enamel phenotype in KTZSL is distinct and more severe than in KS. We propose that SATB1 plays a more stage-specific role during early amelogenesis, as evidenced by the severely hypoplastic enamel in KTZSL patients. In contrast, KMT2D impacts amelogenesis across all stages, as indicated by the hypoplastic and hypomineralized enamel phenotypes in the *Kmt2d*-cKO model. This distinction highlights the broader regulatory effects of KMT2D in coordinating ameloblast differentiation and function throughout enamel formation.

SP6, implicated in hypoplastic amelogenesis imperfecta in humans, is expressed in both presecretory- and secretory-stage ameloblasts. SP6 regulates ameloblast proliferation and differentiation (Bloch-Zupan et al. 2023; Nakamura et al. 2004). Mice deficient in *Sp6* (*Epfn^−/−^*) show enamel deficiencies, as well as defects in cusp and root formation and abnormal dentin structure (Nakamura et al. 2008). Recent studies identified SP6 as a master regulator controlling the expression of *Amelx*, *Ambn*, and *Enam* during tooth development (Rhodes et al. 2021). These studies suggest that SP6 may be a key mediator in KMT2D-regulated early-stage amelogenesis, making it a promising candidate for further investigation.

Among the other candidate genes, *Gpr68*, *Itgb6*, and *Gja1* have been studied in mouse models. Deficiencies in *Gpr68* (*Ogr1*^−/−^) do not result in significant enamel defects, whereas deficiencies in either *Itgb6* (*Itgb6*^−/−^) or *Gja1* (*Gja1^Jrt/+^*) lead to enamel hypoplasia (Flenniken et al. 2005; Mohazab et al. 2013; Parry et al. 2016; Wazen et al. 2015). ITGB6, a cell surface adhesion receptor that mediates cell-cell and cell-extracellular matrix interactions, is expressed in both secretory and maturation-stage ameloblasts (Mohazab et al. 2013; Wang et al. 2014); *Itgb6*^−/−^ mice exhibit a hypomineralized amelogenesis imperfecta phenotype (Mohazab et al. 2013). GJA1, a gap junction protein, is expressed in both secretory- and maturation-stage ameloblasts, and mice with heterozygous *Gja1* mutation (*Gja1*^Jrt/+^) show hypoplastic enamel defects (Toth et al. 2010). We speculate that *Itgb6* and *Gja1* may mediate KMT2D-regulated amelogenesis, contributing specifically to the maturation-stage ameloblasts and SI/SR-mediated secretory-stage ameloblast functions, respectively.

Our findings align with previous research on histone-modifying epigenetic factors in tooth development (Deng et al. 2020; Kobayashi et al. 2021; Li et al. 2021; Takagiwa et al. 2024; Yu et al. 2022; Zhang et al. 2023; Zhu et al. 2024). However, our study is the first to specifically implicate KMT2D in the regulation of amelogenesis in an *in vivo* model. Interestingly, while *KMT2D* deficiency was reported to disturbs cell cycle activity in LS8 dental epithelial cells (Pang et al. 2021), we observed increased proliferation in the *Kmt2d*-cKO tooth germ. This discrepancy may arise from differences between *in vitro* cell line systems and *in vivo* models, as the latter better recapitulates the complex microenvironment and signaling interactions during development.

This study advances our understanding of the molecular mechanisms underlying amelogenesis and suggests potential epigenetic therapeutic targets for amelogenesis imperfecta. While our research provides significant molecular insights into KMT2D-regulated gene programs that control ameloblast differentiation during early-stage amelogenesis, it would be valuable to explore whether these mechanisms also function during late-stage amelogenesis. The recent development of new Cre driver lines (e.g., *Ambn-Cre*, *Amelx-Cre,* and *Odam-Cre*) through the NIH NIDCR FaceBase initiative (Samuels et al. 2020) offers an opportunity to investigate KMT2D’s role across different enamel formation stages. Future studies utilizing stage-specific Cre-driver models will further delineate KMT2D’s functions. Additionally, examining potential interactions between KMT2D and other histone modifiers during enamel formation and similar mechanisms in other mineralized tissues will provide broader insights into epigenetic regulation in development.

## Author Contributions

J.-M. Lee, H.-J.E. Kwon, contributed to conception, design, data acquisition and interpretation, performed statistical analyses, drafted, and critically revised the manuscript. H. Jung, Q. Tang, L. Li contributed to data acquisition and interpretation, performed statistical analyses, and critically revised the manuscript. S.-K. Lee, J.W. Lee, contributed to conception, design, and critically revised the manuscript. Y. Park, contributed to data interpretation, performed statistical analyses, and critically revised the manuscript. All authors gave final approval and agreed to be accountable for all aspects of the work.

## Acknowledgements

We acknowledge the assistance of Woojung An (University at Buffalo); Victor Yuan (University at Buffalo); Andrew McCall, PhD (University at Buffalo); and Peter Bush (University at Buffalo), in various aspects of the current study. Micro-CT and light microscopy data in this study were acquired at the Optical Imaging and Analysis Facility, School of Dental Medicine, State University of New York at Buffalo. SEM data were acquired at the South Campus Instrument Center, School of Dental Medicine, State University of New York at Buffalo. Finally, we thank the members of the Kwon lab and our colleagues in the Department of Oral Biology for their helpful comments and feedback.

## SUPPLEMENTAL APPENDIX

### Supplemental Materials and Methods

#### Micro-Computed Tomography (CT) Scanning and 3-Dimensional (3D) Reconstruction

The heads of 2- and 10-week-old mice were fixed overnight in 10% formalin and then rinsed with phosphate-buffered saline (PBS). Micro-CT scanning was carried out using a SCANCO µCT 100 system (SCANCO Medical, Zurich, Switzerland) with a nominal resolution of 10 µm. The obtained scans were calibrated to milligrams of hydroxyapatite per cubic centimeter (mg HA/cm³) to ensure precise density measurements. Each image was manually adjusted for rotation and alignment along all axes using FIJI software (Schindelin et al. 2012). Image registration was initially performed manually, followed by refinement using the block matching algorithm in the Fijiyama plugin for precise alignment (Fernandez and Moisy 2021). To reduce background noise, areas with values below 200.03 mg HA/cm³ were excluded. Images were then cropped to isolate the specific region of interest within the first molars and incisors. For the generation of representative frontal section planes along the molar and incisor, the Multi Kymograph program in the FIJI plugin was utilized. This program facilitated consistent placement of the section planes at defined anatomical landmarks. Finally, 3D rendering and visualization were accomplished using the 3Dscript plugin (Schmid et al. 2019). For molars, we ensured consistency in using comparable sections for each section plane by carefully selecting sections based on identifiable landmarks, specifically the tallest point of the mesiobuccal ‘protoconid’ and ‘metaconid’ cusps of the first molar. This section level consistently transects through these cusp tips to enable reproducible comparisons across samples.

#### Scanning Electron Microscopy (SEM) Imaging

Mouse mandibles were collected and fixed in 4% paraformaldehyde in 10 mM PBS at room temperature for 1 hour. After fixation, the tissues were stored overnight at 4°C. To ensure the complete removal of any residual fixative, the tissues were thoroughly washed with PBS. The gingival tissues of the mandibles were then carefully dissected to expose the teeth and surrounding periodontal structures. Importantly, decalcification was omitted to preserve the bone tissue in its natural mineralized state. The enamel was etched with 37.5% KERR Gel etchant. After etching, the specimens were rinsed with phosphoric acid and then thoroughly washed with water to remove any remaining etchant. The specimens were mounted and coated with evaporated carbon using a high vacuum evaporator (Denton 502 Evaporator, Denton Vacuum, Moorestown, NJ, USA). SEM was performed using a Hitachi SU-70 field emission SEM (Tokyo, Japan) at an accelerating voltage of 2.0 keV. Imaging was conducted using an in-lens secondary electron detector at zero tilt, or with a lower detector at a 70° tilt, to achieve detailed visualization of the molar surface structures.

#### Histology and Immunofluorescence Staining

Mice were euthanized at designated developmental stages. Head tissue specimens were fixed in 4% paraformaldehyde, dehydrated through a graded ethanol series, embedded in paraffin, and sectioned at a thickness of 5–7 μm, as previously reported (Lee et al. 2022). Specimens collected at E16.5 and P0.5 were decalcified in 14% ethylenediaminetetraacetic acid (EDTA) for 3 and 7 days, respectively. For histological analysis, sections were stained with hematoxylin and eosin, as previously described (Lee et al. 2022), and imaged using an Axioscope upright light microscope at 20X magnification (Zeiss, Oberkochen, Germany). An SZX16 stereomicroscope (Olympus, Tokyo, Japan) was used for capturing gross images. For immunofluorescence staining, antigen retrieval was conducted in an antigen retrieval buffer using a pressure cooker for 15 minutes at low pressure. Sections were then washed in PBS and blocked at room temperature in a blocking buffer containing 2% goat serum, 5% BSA, 1% Triton-X 100, and 0.1% Tween-20 in PBS for 1 hour. Primary antibodies, diluted in blocking buffer, were incubated with the sections overnight at 4°C. The following day, sections were washed in PBS, incubated with secondary antibodies diluted in blocking buffer for 1 hour at room temperature, washed again in PBS, and mounted using Vectashield mounting media with DAPI (Vector Laboratories, Burlingame, CA, USA; H-1200). The primary antibody used was a homemade guinea pig anti-KMT2D (1:300) (Huisman et al. 2021). Immunofluorescence images were captured using a THUNDER Imager fluorescence microscope (Leica Microsystems, Wetzlar, Germany).

#### EdU Staining

EdU (5-ethynyl-2’-deoxyuridine) was administered to pregnant mice at a dosage of 10 mg/kg of body weight, one hour before the mice were euthanized. The embryos were then dissected, processed, and sectioned at a thickness of 5 μm. EdU detection was performed using the Click-iT Plus Alexa Fluor 488 imaging kit according to the manufacturer’s instructions (Invitrogen, Carlsbad, CA, USA; C10637).

#### TUNEL Assay

The DeadEnd™ Fluorometric TUNEL System (Promega, Madison, WI, USA; G3250) was used to assess apoptosis in paraffin sections, according to the manufacturer’s protocol.

#### Skeletal Preparations

Mouse heads, collected at P0.5, were fixed in 100% ethanol for 3 days, stained with an Alcian Blue solution (30 mg Alcian Blue dissolved in 20 ml glacial acetic acid and 80 ml 95% ethanol) for 2 days, and then re-fixed in 100% ethanol for 24 hours. The samples were subsequently cleared in 2% KOH for 3 hours and counterstained with an Alizarin Red solution (50 mg Alizarin Red dissolved in 1 L of 1% KOH) for 24 hours. Following staining, tissue clearing was performed in 1% KOH/20% glycerol for 3 days. Images were captured using a standard stereomicroscope (Amscope, Irvine, CA, USA).

#### RNA-Seq and Bioinformatics Analysis

Total RNA was isolated from micro-dissected first molars of control and *Kmt2d* conditional knockout (cKO) mice at birth (P0.5) using the PureLink RNA Mini Kit (Invitrogen, Carlsbad, CA, USA; 12183018A). Four biological replicates per genotype, control and *Kmt2d*-cKO, were utilized for sequencing. Library preparation and high-throughput sequencing (Illumina, San Diego, CA, USA; NovaSeq 6000) were carried out by Novogene (Beijing, China). Sequencing reads were processed using Salmon (Patro et al. 2017) to quantify transcript-level gene expression. The resulting Salmon outputs were further analyzed using DESeq2 (Love et al. 2014) to assess gene expression at the gene level and identify differentially expressed genes (DEGs) between *Kmt2d*-cKO and littermate controls. DEGs were defined by a log_2_ fold change (log_2_FC) ≥ 0.4 and an adjusted p-value ≤ 0.001. Volcano plots were generated using the EnhancedVolcano package (Blighe et al. 2024). Gene ontology analysis was conducted with the clusterProfiler package (Wu et al. 2021). The RNA-seq data files have been submitted to the National Center for Biotechnology Information Gene Expression Omnibus (NCBI GEO) database under BioProject accession number PRJNA1149088.

#### CUT&RUN-Seq and CUT&RUN-qPCR

Cells were isolated from the first molars of wildtype mice at birth (P0.5), with approximately 15,000 cells per group utilized for the CUT&RUN assay, following the manufacturer’s protocol (Cell Signaling, Danvers, MA, USA; 86652). Library preparation was performed using the NEBNext UltraII DNA Library Prep Kit for Illumina (New England Biolabs, Ipswich, MA, USA; NEB#7645L) as per the provided instructions. Sequencing reads were mapped to the mouse genome (mm10) by Bowtie 2 (Langmead and Salzberg 2012). The mapping rates were 94.52% and 62.11% for the first and second replicates, respectively. The mapped reads were then analyzed by MACS3 (Zhang et al. 2008) to identify KMT2D binding peaks. For the two replicates, we identified 4,198 and 3,030 peaks, respectively. BEDTools (Quinlan and Hall 2010) identified 2,271 peaks that are common to both replicates. We used the GREAT webserver (McLean et al. 2010) for the overrepresentation analysis of gene ontology terms for the 2,271 KMT2D binding peaks. The CUT&RUN-seq data files have been deposited in the National Center for Biotechnology Information Gene Expression Omnibus (NCBI GEO) database under BioProject accession number PRJNA1149088.

Specific genomic loci were quantified by qPCR using primers designed to target regions of interest. *Untr6* was used as reference. Relative fold enrichment at each locus was calculated using the ΔΔCt method (Panday et al. 2022), normalizing to IgG controls.

#### CUT&RUN qPCR Primer Sequences

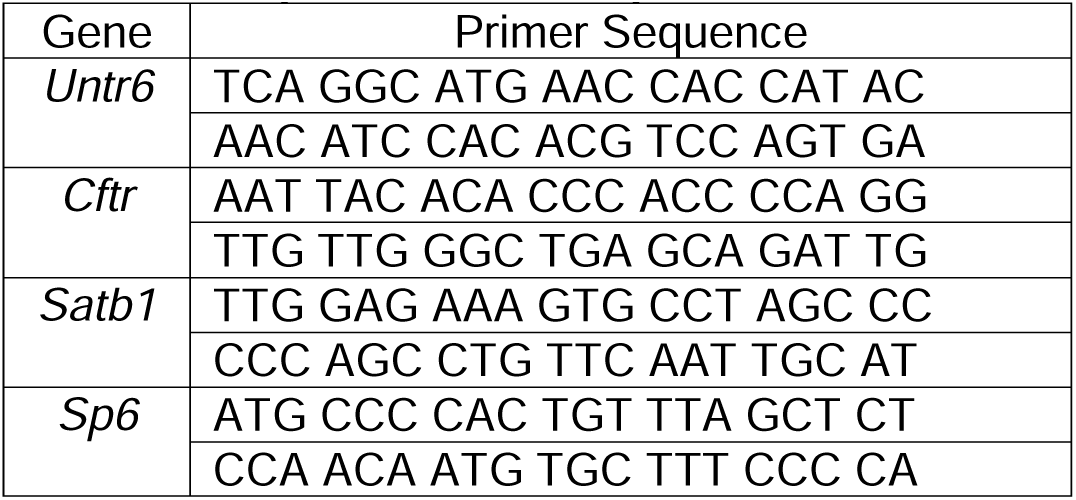

#### Analysis of Publicly Available Single-cell RNA Sequencing (scRNA-Seq) Datasets

The scRNA-seq datasets performed on mouse P7 incisors (GSE146855) were imported into Seurat v3.6 (Stuart et al. 2019) for unsupervised clustering and cell population classification. Subsequently, dental epithelial cells were subsetted and sub-clustered to identify major dental epithelial cell types, which were then projected onto a UMAP (Uniform Manifold Approximation and Projection) plot. The expression of *Kmt2d*/*Mll4* in major dental epithelial cell types was visualized with FeaturePlot function implemented in the Seurat package. The joint expression of *Kmt2d* and its direct target genes were estimated and visualized using the Nebulosa package (Alquicira-Hernandez and Powell 2021).

#### Dispase Treatment for Dental Epithelium and Mesenchyme Separation

Mandibular first molars were dissected from control and *Kmt2d* conditional knockout (cKO) mice at birth (P0.5) under a stereomicroscope. The dissected molars were incubated in 2.2 U/mL dispase II (Sigma-Aldrich) dissolved in PBS at 37°C for 30 minutes. Following enzymatic digestion, the dental epithelium and mesenchyme were carefully separated using fine-tipped forceps under a stereomicroscope. The isolated tissues were then processed for RNA extraction using the RNeasy Plus Micro Kit (Qiagen) or for the CUT&RUN assay, both performed according to the manufacturer’s instructions.

#### Quantitative Real-Time RT-PCR (qRT-PCR)

Total RNA was reverse-transcribed with iScript™ Reverse Transcription Supermix (Bio-Rad, Hercules, CA, USA). qRT-PCR was conducted using the 7500 Real-Time PCR System (Applied Biosystems, Foster City, CA, USA) and SsoAdvanced Universal SYBR Green Supermix (Bio-Rad, Hercules, CA, USA). The expression levels of the target genes were normalized to *Hprt* expression. PCR reactions were performed in triplicate. Differential expression was analyzed using Student’s *t*-test.

#### qRT-PCR Primer Sequences

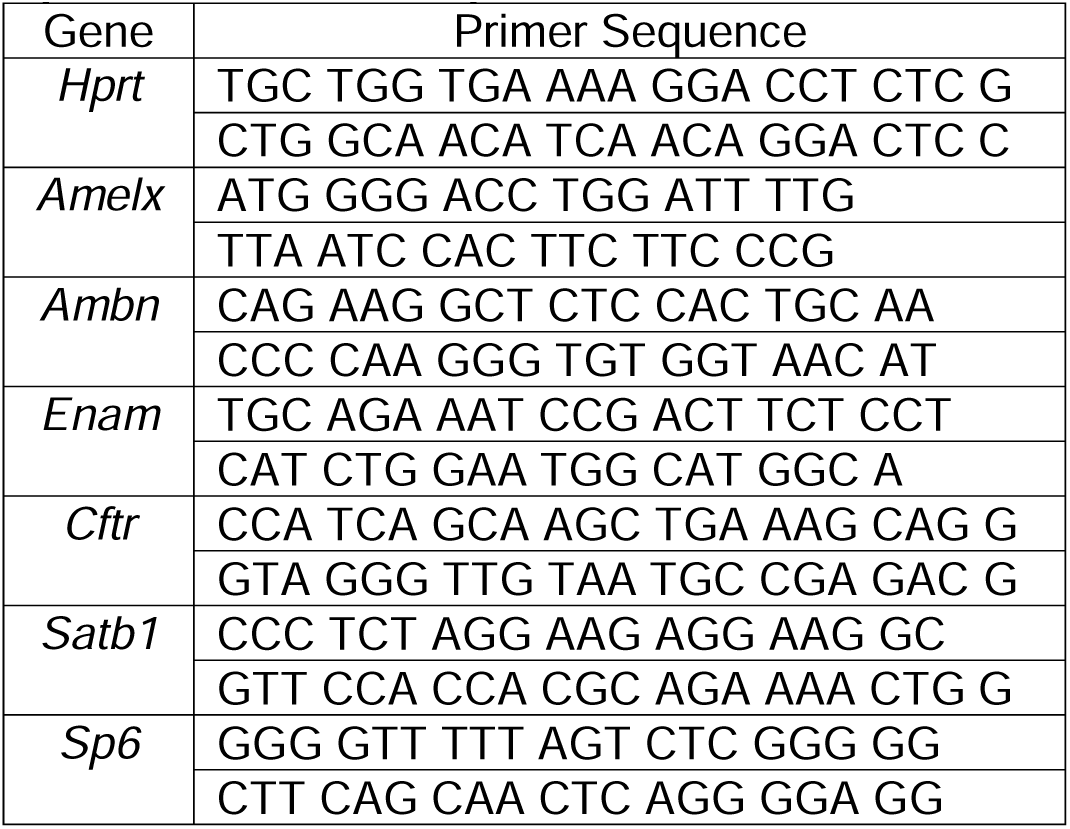

#### Statistics

All statistical analyses were conducted using Prism 10 software (GraphPad Software, San Diego, CA, USA). Pairwise comparisons for differential expression were assessed using Student’s *t*-test. Statistical significance was determined with *p*-values defined as ≤ 0.05 (*), ≤ 0.01 (**), ≤ 0.001 (***), and ≤ 0.0001 (****).

### Supplemental Figures

**Appendix Figure 1.**
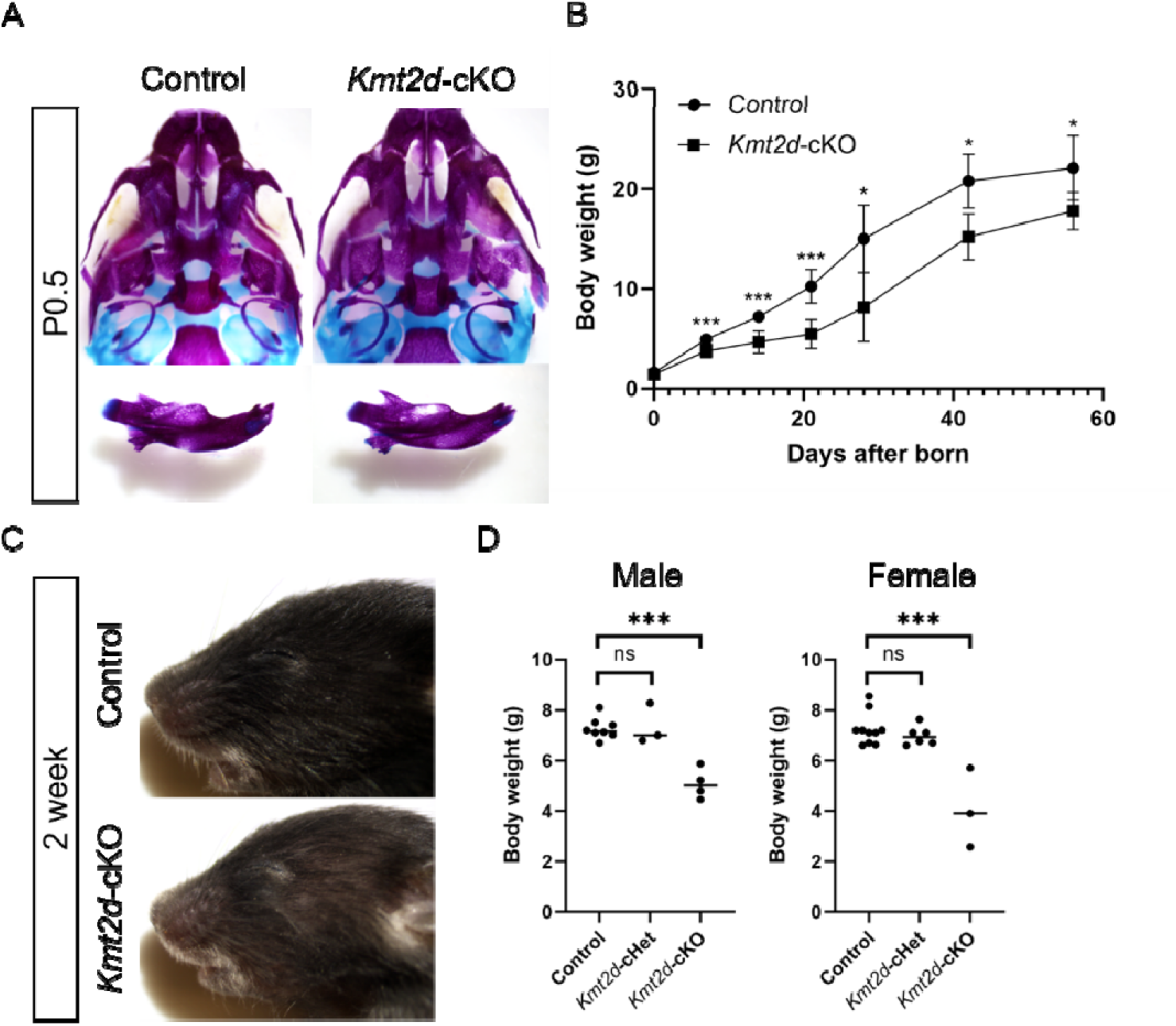
*Kmt2d*-cKO mice do not show visible craniofacial anomalies but exhibit hair thinning and reduced body weight. (A) Skeletal staining of the maxillary (top row) and mandibular jaws (bottom row). Overall skeletal structures were similar between control and *Kmt2d*-cKO mice at birth (P0.5). (B) Body weight kinetics showing significantly reduced body weight in the *Kmt2d*-cKO mice compared to controls at all postnatal time points. (C) Lateral view of the craniofacial region at 2 weeks, showing scattered hair thinning but no visible craniofacial differences between control and *Kmt2d*-cKO mice. (D) Body weights of control, *Kmt2d*-cHet, and *Kmt2d*-cKO mice, both male and female, at 2 weeks. No significant difference was found between control and *Kmt2d*-cHet mice. However, *Kmt2d*-cKO mice showed significantly reduced body weight compared to controls. \**p* < 0.05; \*\*\**p* < 0.001; ns, *p* > 0.05. *n* = 3–10 per group.

**Appendix Figure 2.**
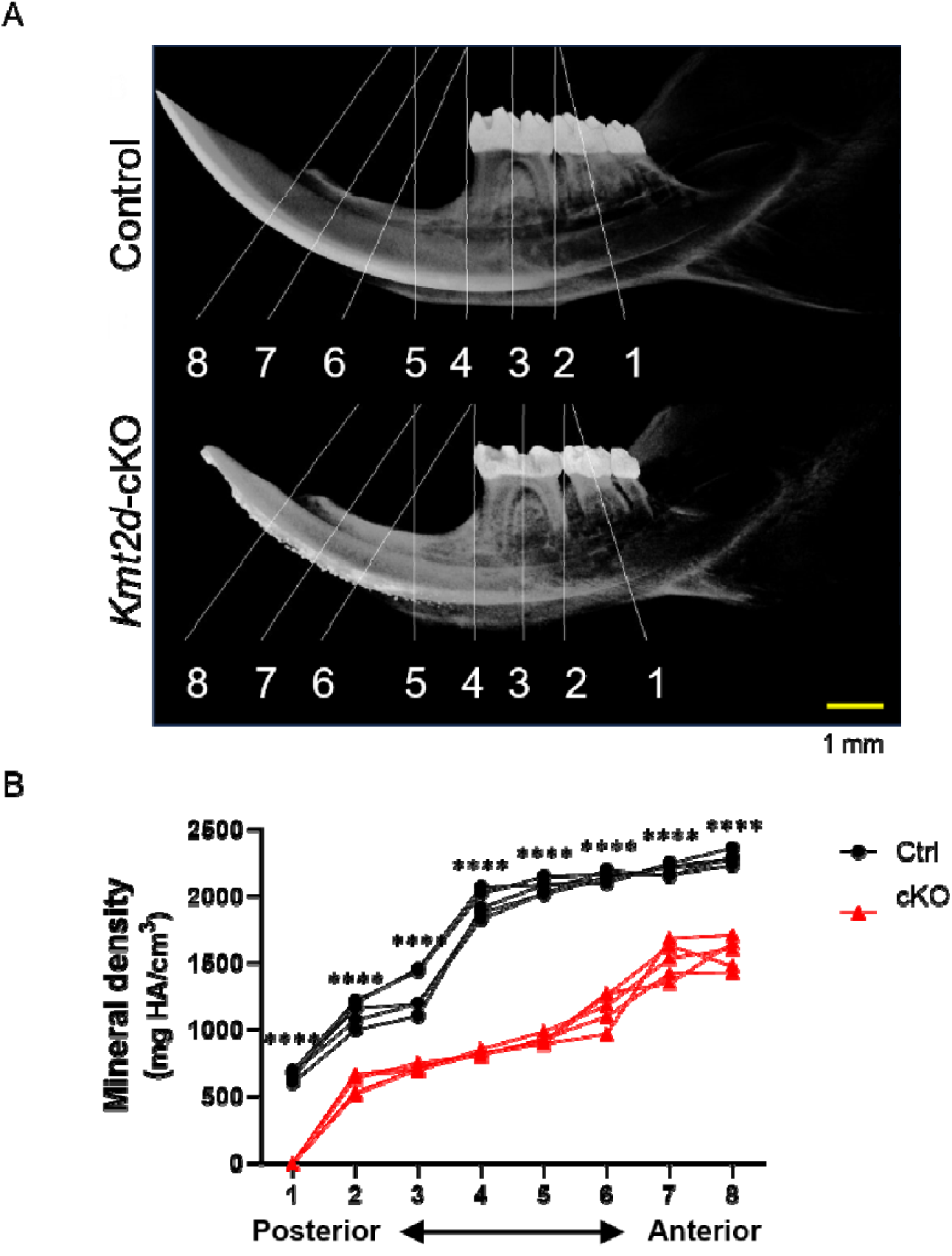
Defective enamel in the *Kmt2d*-cKO mouse incisor. (A, B) Micro-CT-based comparison of control and *Kmt2d*-cKO mouse incisors at 8 weeks. (A) 3D volume-rendered images, or pseudo-X-ray views, show eight representative section planes along the anteroposterior axis of the mandible, based on skeletal and dental anatomical landmarks (Bui et al. 2023; Hu et al. 2011). These planes correspond to the upper part of the incisor exposure point (8), the midsection between levels 8 and 6 (7), the lower part of the incisor exposure point (6), the minimum depression on the dorsal side of the incisor ramus to the anterior margin of the muscle insertion area on the ventral side of the incisor ramus (5) (Boell et al. 2013), the mesial section of M1 (4), the midsection of M1 (3), the distal section of M1 (2), and the midsection of M2 (1). Scale bar: 1 mm. (B) Mineral density distribution along the anteroposterior axis of incisors in 8-week-old mice. Mineral density (mg HA/cm³) was measured at the eight section planes described above. Control (Ctrl): black circles; *Kmt2d*-cKO (cKO): red triangles. \*\*\*\**p* < 0.0001; *n* = 5–6 per group.

**Appendix Figure 3.**
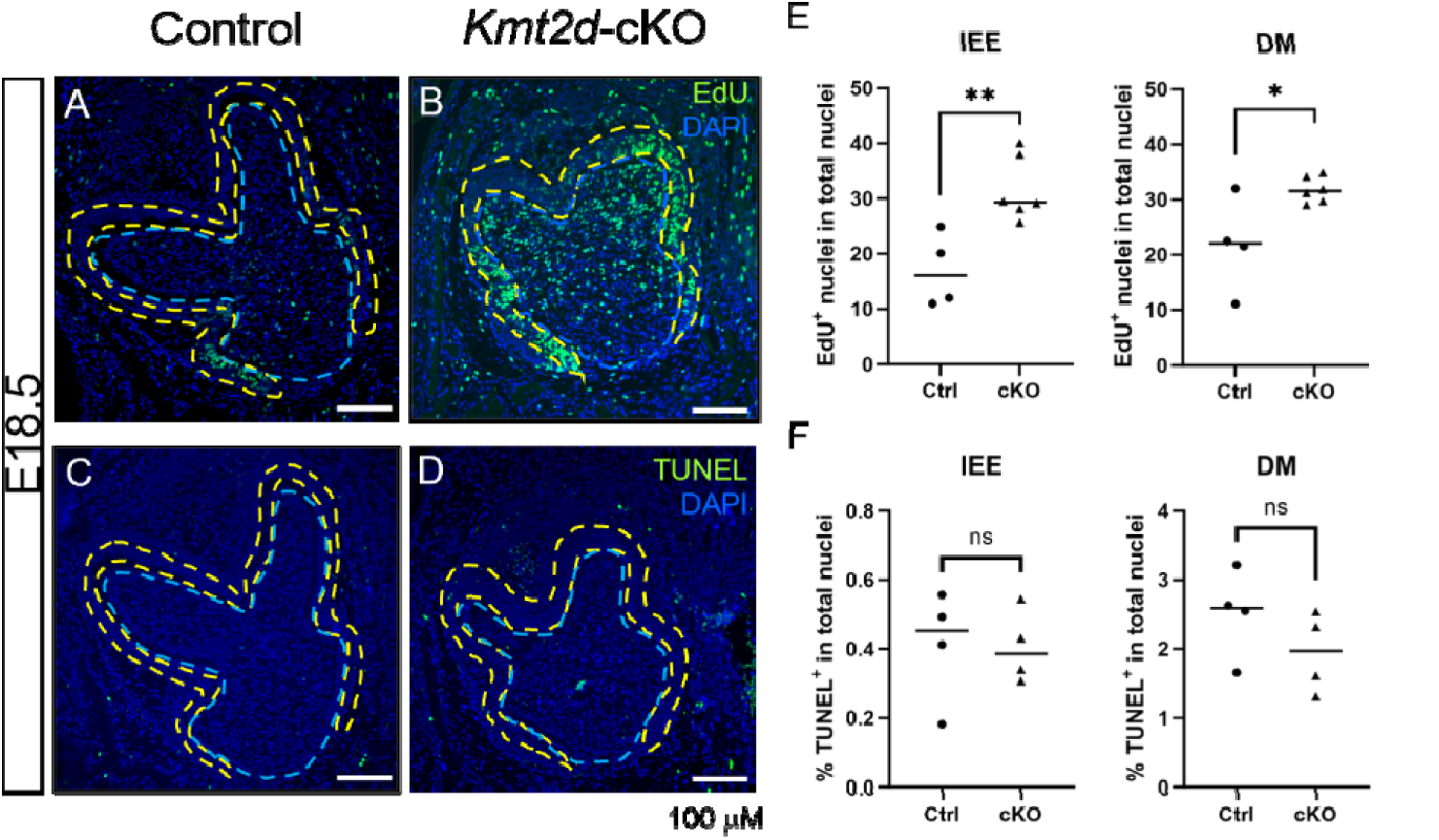
*Kmt2d*-cKO mouse embryos exhibit increased proliferation in both the pre-ameloblast and pre-odontoblast layers, while apoptosis remain unchanged. (A, B, E) EdU staining showed increased proliferation pre-ameloblast (dental epithelial) and pre-odontoblast (dental mesenchymal) layers of the *Kmt2d*-cKO first molars compared to controls at E18.5. (C, D, F) TUNEL staining revealed no significant changes in apoptosis in either layer. PreAmB, pre-ameloblast (outlined with yellow dashed lines); PreOd, pre-odontoblast (outlined with light blue dashed lines). \**p* < 0.05; \*\**p* < 0.01; ns, *p* > 0.05. *n* = 4–6 per group.

**Appendix Figure 4.**
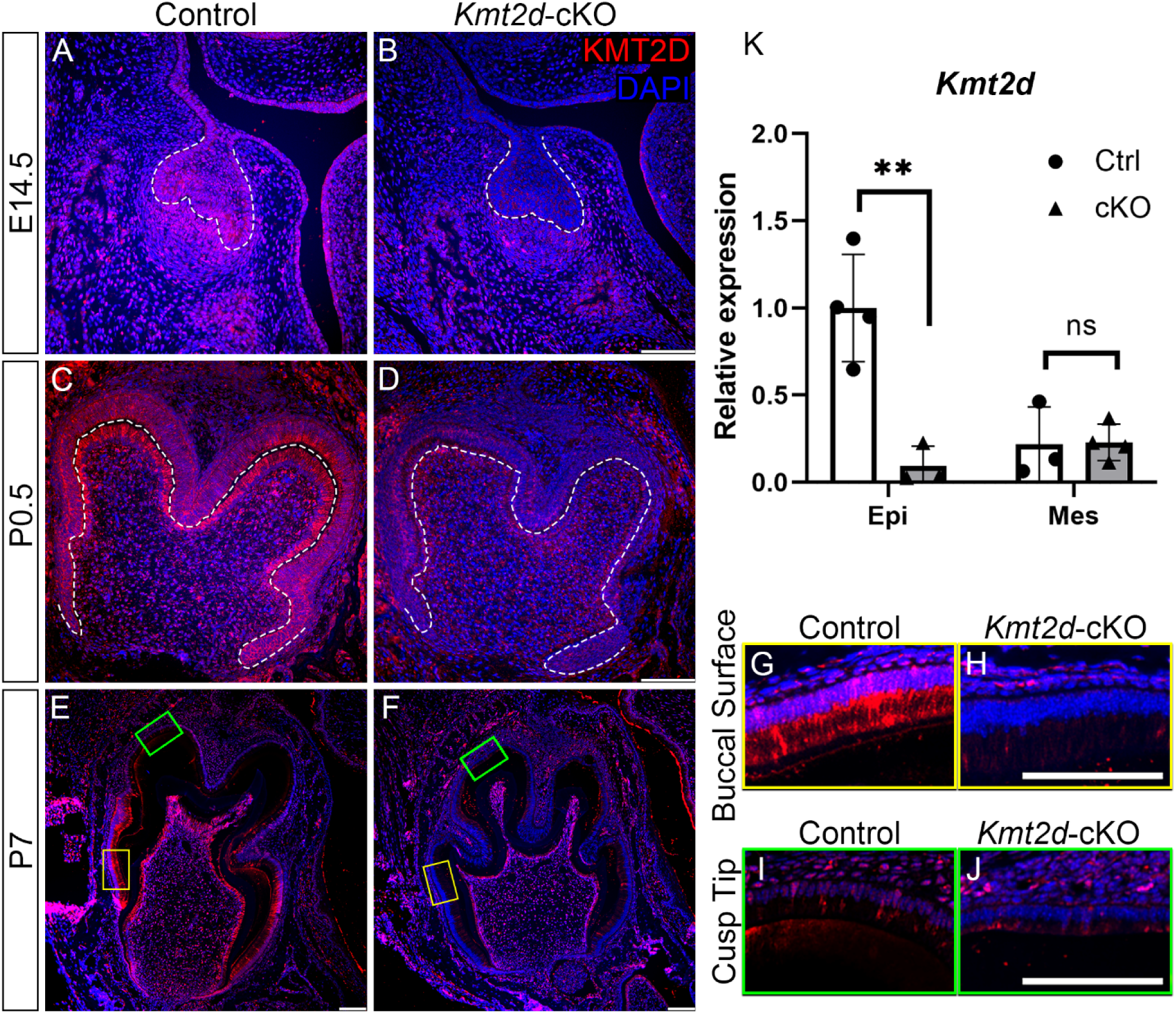
*Kmt2d*-cKO tooth germs exhibit decreased KMT2D expression in the dental epithelium. (A–J) Immunofluorescence staining for KMT2D in the first molar of control and *Kmt2d*-cKO mice at E14.5 (A, B), P0.5 (C, D), and P7 (E–J), showing markedly reduced KMT2D signals in the dental epithelium as well as dental mesenchyme of *Kmt2d*-cKO (B, D, F, H, J) compared to control (A, C, E, G, I) tooth germs. The dental epithelium-mesenchyme interface is outlined with white dashed lines (in panels A–D). Higher magnifications of the buccal surface (G, H; yellow boxes in panels E, F) and the cusp tip (I, J; green boxes in panels E, F) of the mesiobuccal ‘protoconid’ cusp both show reduced KMT2D signals in the ameloblast layer in the *Kmt2d*-cKO mice compared to controls. *n* = 4 per group. (K) qRT-PCR showing significantly decreased epithelial *Kmt2d* expression (Epi), while mesenchymal *Kmt2d* expression (Mes) remained unaffected in the *Kmt2d*-cKO first molar compared to the control. \**p* < 0.05. *n* = 3–4 per group.

**Appendix Figure 5.**
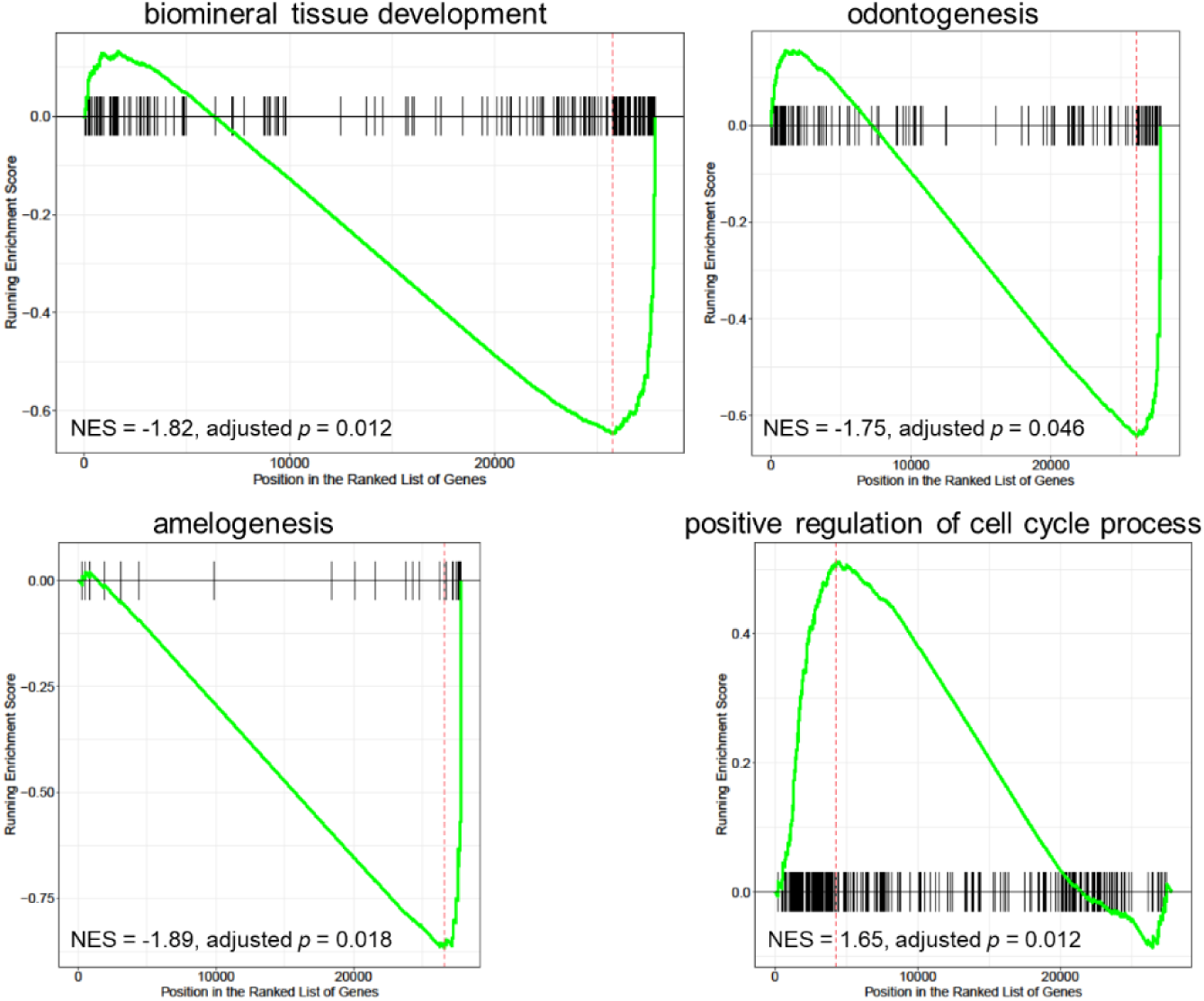
Gene set enrichment (GSE) analysis-based enriched biological processes in *Kmt2d*-cKO first molars at P0.5.

**Appendix Figure 6.**
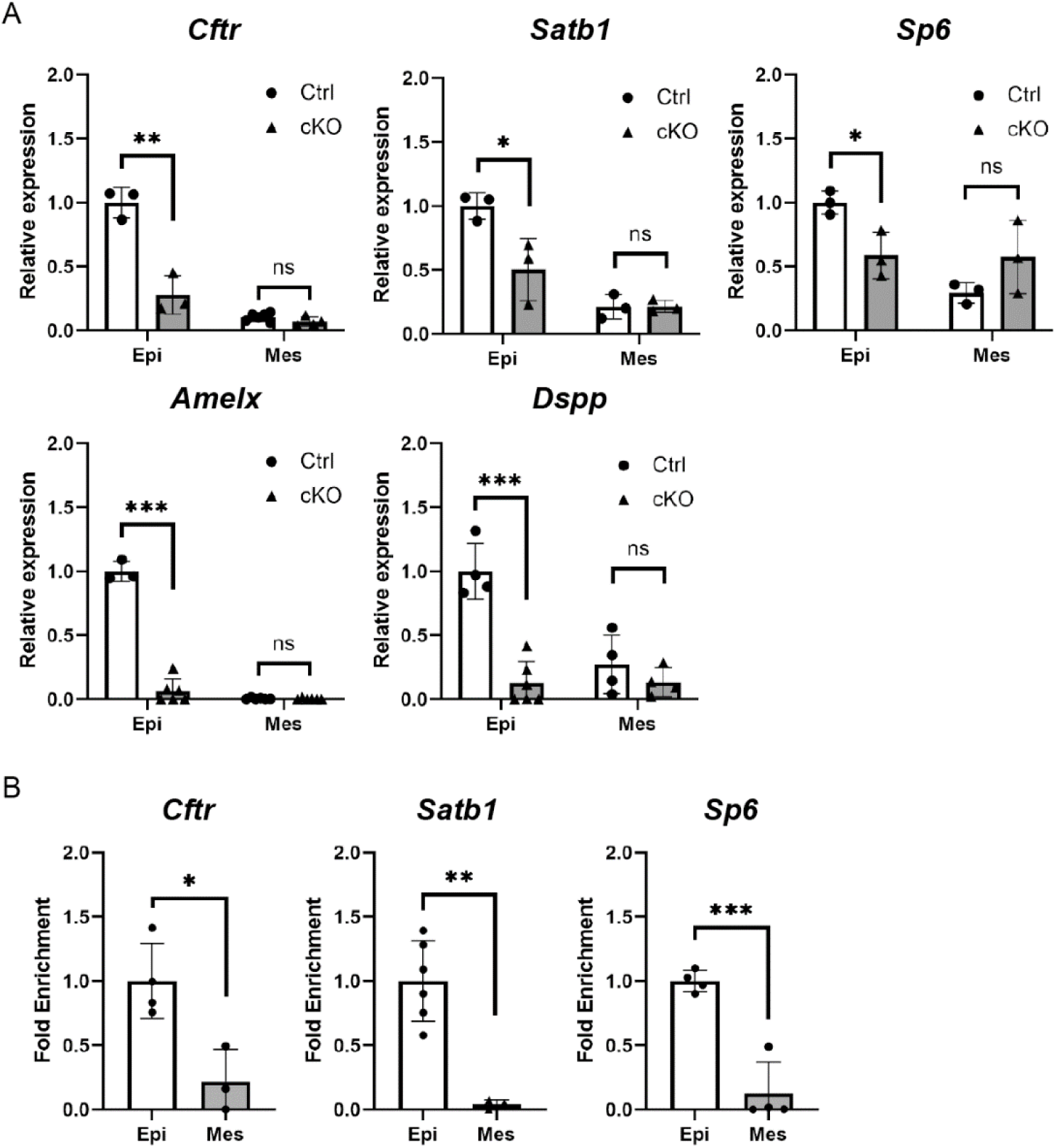
Tissue-specific mRNA expression changes and fold enrichment of amelogenesis-related genes. (A) qRT-PCR analysis of Cftr, Satb1, Sp6, Amelx, and Dspp, showing successful reduction of their mRNA expression exclusively in the Kmt2d-cKO dental epithelium, with no changes observed in the Kmt2d-cKO dental mesenchyme. (B) CUT&RUN-qPCR analysis showing specific fold enrichment of Cftr, Satb1, and Sp6 in dental epithelium compared to the dental mesenchyme. *p < 0.05; **p < 0.01; ***p < 0.001; ns, p > 0.05. n = 3–6 per group.

**Appendix Table 1.**
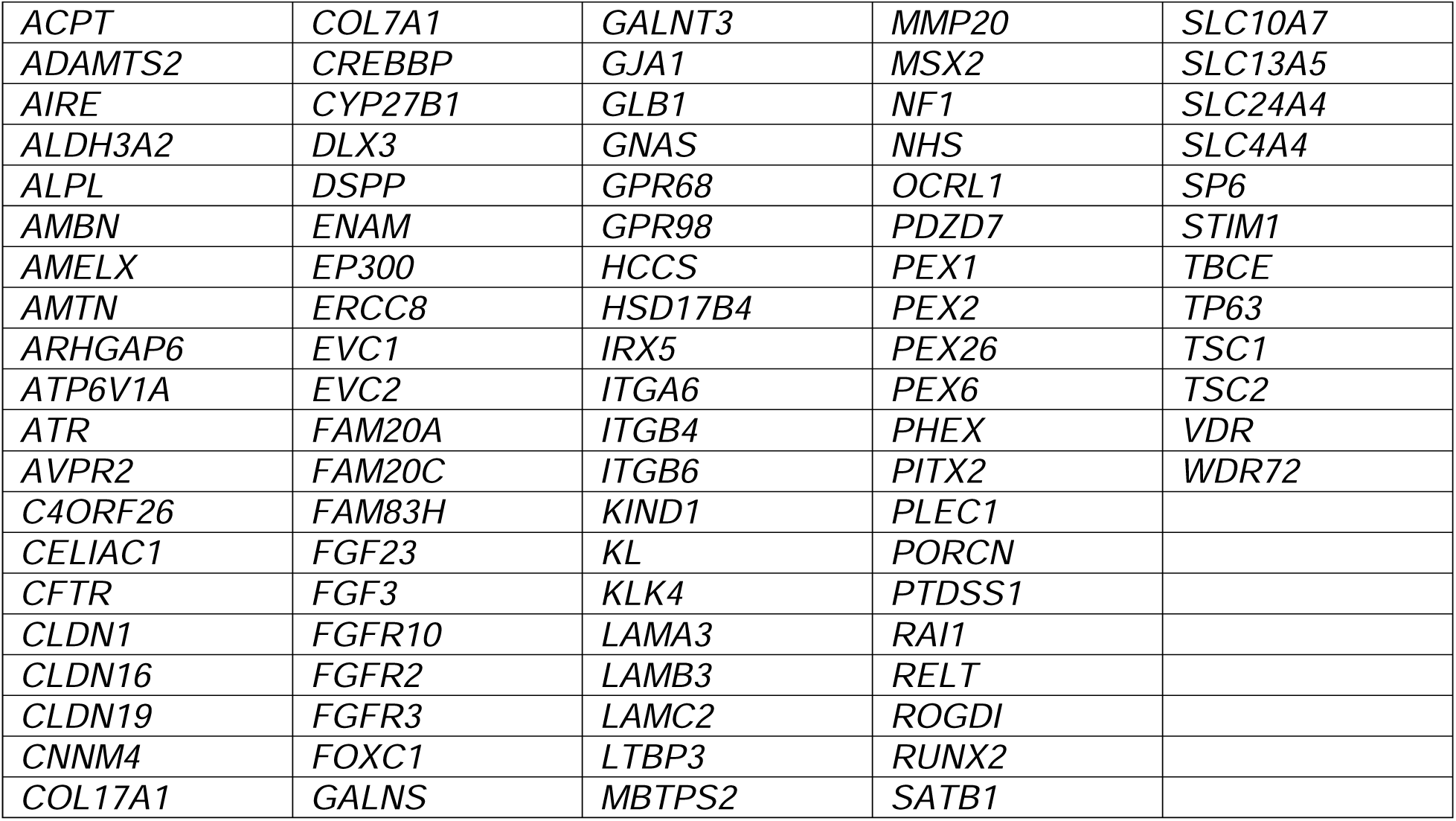
List of the 92 amelogenesis-related genes. A total of 92 genes were compiled from previous studies (Bloch-Zupan et al. 2023; Dong et al. 2023; Smith et al. 2017; Wright et al. 2015).

