## Supplemental Methods, Appendix Figures for "KMT2D regulates tooth enamel development"

***CUT&RUN qPCR Primer Sequences***

| Gene | Primer Sequence |
| --- | --- |
| *Untr6* | TCA GGC ATG AAC CAC CAT AC |
|  | AAC ATC CAC ACG TCC AGT GA |
| *Cftr* | AAT TAC ACA CCC ACC CCA GG |
|  | TTG TTG GGC TGA GCA GAT TG |
| *Satb1* | TTG GAG AAA GTG CCT AGC CC |
|  | CCC AGC CTG TTC AAT TGC AT |
| *Sp6* | ATG CCC CAC TGT TTA GCT CT |
|  | CCA ACA ATG TGC TTT CCC CA |

***Analysis of Publicly Available Single-cell RNA Sequencing (scRNA-Seq) Datasets***

The scRNA-seq datasets performed on mouse P7 incisors (GSE146855) were imported into Seurat v3.6 (Stuart et al. 2019) for unsupervised clustering and cell population classification. Subsequently, dental epithelial cells were subsetted and sub-clustered to identify major dental epithelial cell types, which were then projected onto a UMAP (Uniform Manifold Approximation and Projection) plot. The expression of *Kmt2d*/*Mll4* in major dental epithelial cell types was visualized with FeaturePlot function implemented in the Seurat package. The joint expression of *Kmt2d* and its direct target genes were estimated and visualized using the Nebulosa package (Alquicira-Hernandez and Powell 2021).

***Dispase Treatment for Dental Epithelium and Mesenchyme Separation***

Mandibular first molars were dissected from control and *Kmt2d* conditional knockout (cKO) mice at birth (P0.5) under a stereomicroscope. The dissected molars were incubated in 2.2 U/mL dispase II (Sigma-Aldrich) dissolved in PBS at 37°C for 30 minutes. Following enzymatic digestion, the dental epithelium and mesenchyme were carefully separated using fine-tipped forceps under a stereomicroscope. The isolated tissues were then processed for RNA extraction using the RNeasy Plus Micro Kit (Qiagen) or for the CUT&RUN assay, both performed according to the manufacturer’s instructions.

***Quantitative Real-Time RT-PCR (qRT-PCR)***

Total RNA was reverse-transcribed with iScript™ Reverse Transcription Supermix (Bio-Rad, Hercules, CA, USA). qRT-PCR was conducted using the 7500 Real-Time PCR System (Applied Biosystems, Foster City, CA, USA) and SsoAdvanced Universal SYBR Green Supermix (Bio-Rad, Hercules, CA, USA). The expression levels of the target genes were normalized to *Hprt* expression. PCR reactions were performed in triplicate. Differential expression was analyzed using Student’s *t*-test.

***qRT-PCR Primer Sequences***

| Gene | Primer Sequence |
| --- | --- |
| *Hprt* | TGC TGG TGA AAA GGA CCT CTC G |
|  | CTG GCA ACA TCA ACA GGA CTC C |
| *Amelx* | ATG GGG ACC TGG ATT TTG |
|  | TTA ATC CAC TTC TTC CCG |
| *Ambn* | CAG AAG GCT CTC CAC TGC AA |
|  | CCC CAA GGG TGT GGT AAC AT |
| *Enam* | TGC AGA AAT CCG ACT TCT CCT |
|  | CAT CTG GAA TGG CAT GGC A |
| *Cftr* | CCA TCA GCA AGC TGA AAG CAG G |
|  | GTA GGG TTG TAA TGC CGA GAC G |
| *Satb1* | CCC TCT AGG AAG AGG AAG GC |
|  | GTT CCA CCA CGC AGA AAA CTG G |
| *Sp6* | GGG GTT TTT AGT CTC GGG GG |
|  | CTT CAG CAA CTC AGG GGA GG |

***Statistics***

All statistical analyses were conducted using Prism 10 software (GraphPad Software, San Diego, CA, USA). Pairwise comparisons for differential expression were assessed using Student’s *t*-test. Statistical significance was determined with *p*-values defined as ≤ 0.05 (*), ≤ 0.01 (**), ≤ 0.001 (***), and ≤ 0.0001 (****).

**Supplemental Figures**

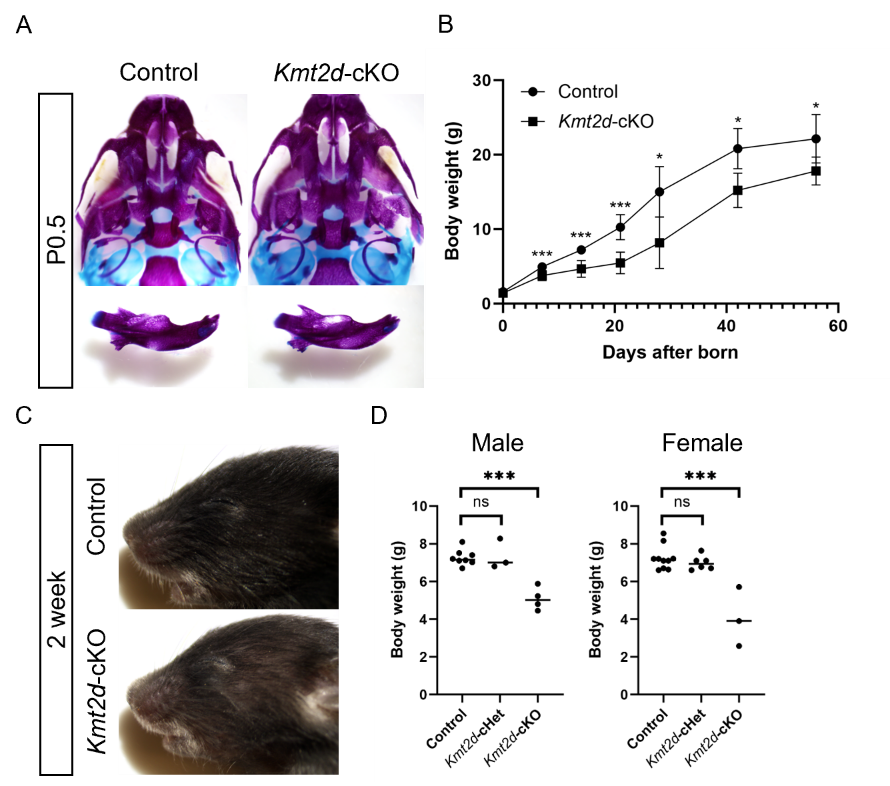

**Appendix Figure 1. *Kmt2d*-cKO mice do not show visible craniofacial anomalies but exhibit hair thinning and reduced body weight.** (A) Skeletal staining of the maxillary (top row) and mandibular jaws (bottom row). Overall skeletal structures were similar between control and *Kmt2d*-cKO mice at birth (P0.5). (B) Body weight kinetics showing significantly reduced body weight in the *Kmt2d*-cKO mice compared to controls at all postnatal time points. (C) Lateral view of the craniofacial region at 2 weeks, showing scattered hair thinning but no visible craniofacial differences between control and *Kmt2d*-cKO mice. (D) Body weights of control, *Kmt2d*-cHet, and *Kmt2d*-cKO mice, both male and female, at 2 weeks. No significant difference was found between control and *Kmt2d*-cHet mice. However, *Kmt2d*-cKO mice showed significantly reduced body weight compared to controls. **p* < 0.05; ****p* < 0.001; ns, *p* > 0.05. *n* = 3–10 per group.

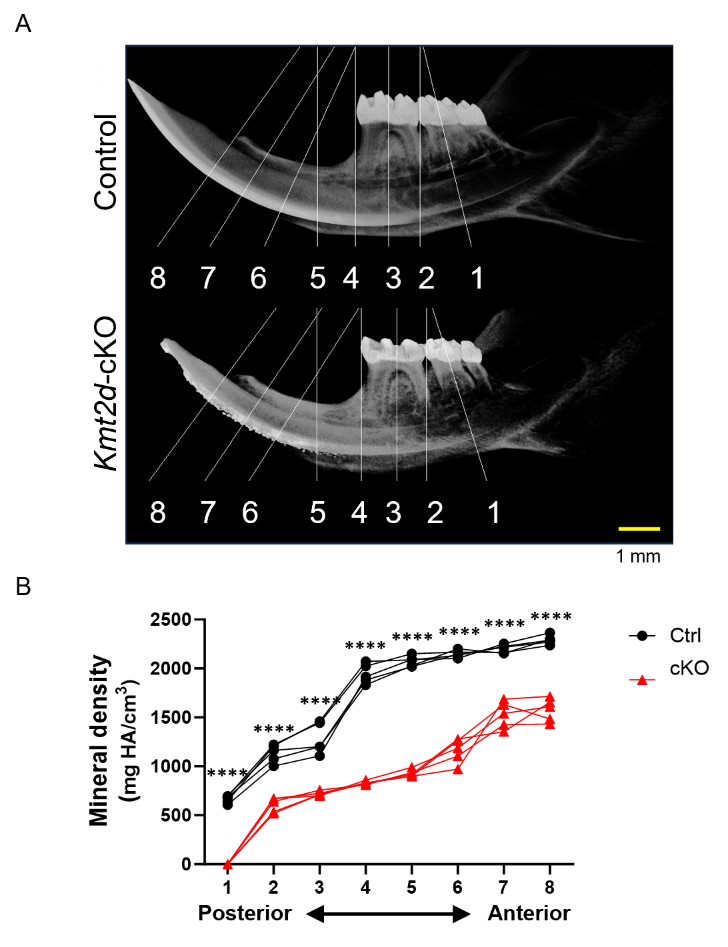

**Appendix Figure 2. Defective enamel in the *Kmt2d*-cKO mouse incisor.** (A, B) Micro-CT-based comparison of control and *Kmt2d*-cKO mouse incisors at 8 weeks. (A) 3D volume-rendered images, or pseudo-X-ray views, show eight representative section planes along the anteroposterior axis of the mandible, based on skeletal and dental anatomical landmarks (Bui et al. 2023; Hu et al. 2011). These planes correspond to the upper part of the incisor exposure point (8), the midsection between levels 8 and 6 (7), the lower part of the incisor exposure point (6), the minimum depression on the dorsal side of the incisor ramus to the anterior margin of the muscle insertion area on the ventral side of the incisor ramus (5) (Boell et al. 2013), the mesial section of M1 (4), the midsection of M1 (3), the distal section of M1 (2), and the midsection of M2 (1). Scale bar: 1 mm. (B) Mineral density distribution along the anteroposterior axis of incisors in 8-week-old mice. Mineral density (mg HA/cm³) was measured at the eight section planes described above. Control (Ctrl): black circles; *Kmt2d*-cKO (cKO): red triangles. *****p* < 0.0001; *n* = 5–6 per group.

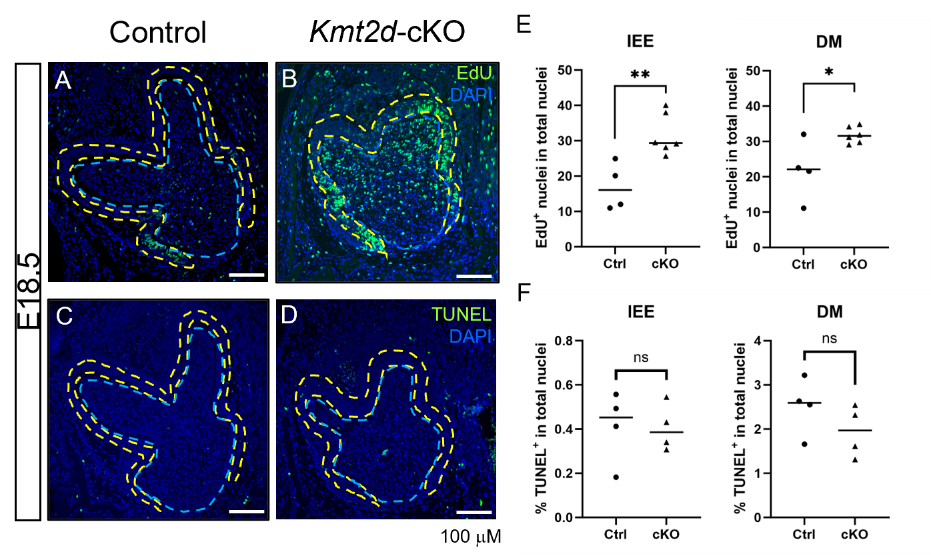

**Appendix Figure 3. *Kmt2d*-cKO mouse embryos exhibit increased proliferation in both the pre-ameloblast and pre-odontoblast layers, while apoptosis remain unchanged.** (A, B, E) EdU staining showed increased proliferation pre-ameloblast (dental epithelial) and pre-odontoblast (dental mesenchymal) layers of the *Kmt2d*-cKO first molars compared to controls at E18.5. (C, D, F) TUNEL staining revealed no significant changes in apoptosis in either layer. PreAmB, pre-ameloblast (outlined with yellow dashed lines); PreOd, pre-odontoblast (outlined with light blue dashed lines). **p* < 0.05; ***p* < 0.01; ns, *p* > 0.05. *n* = 4–6 per group.

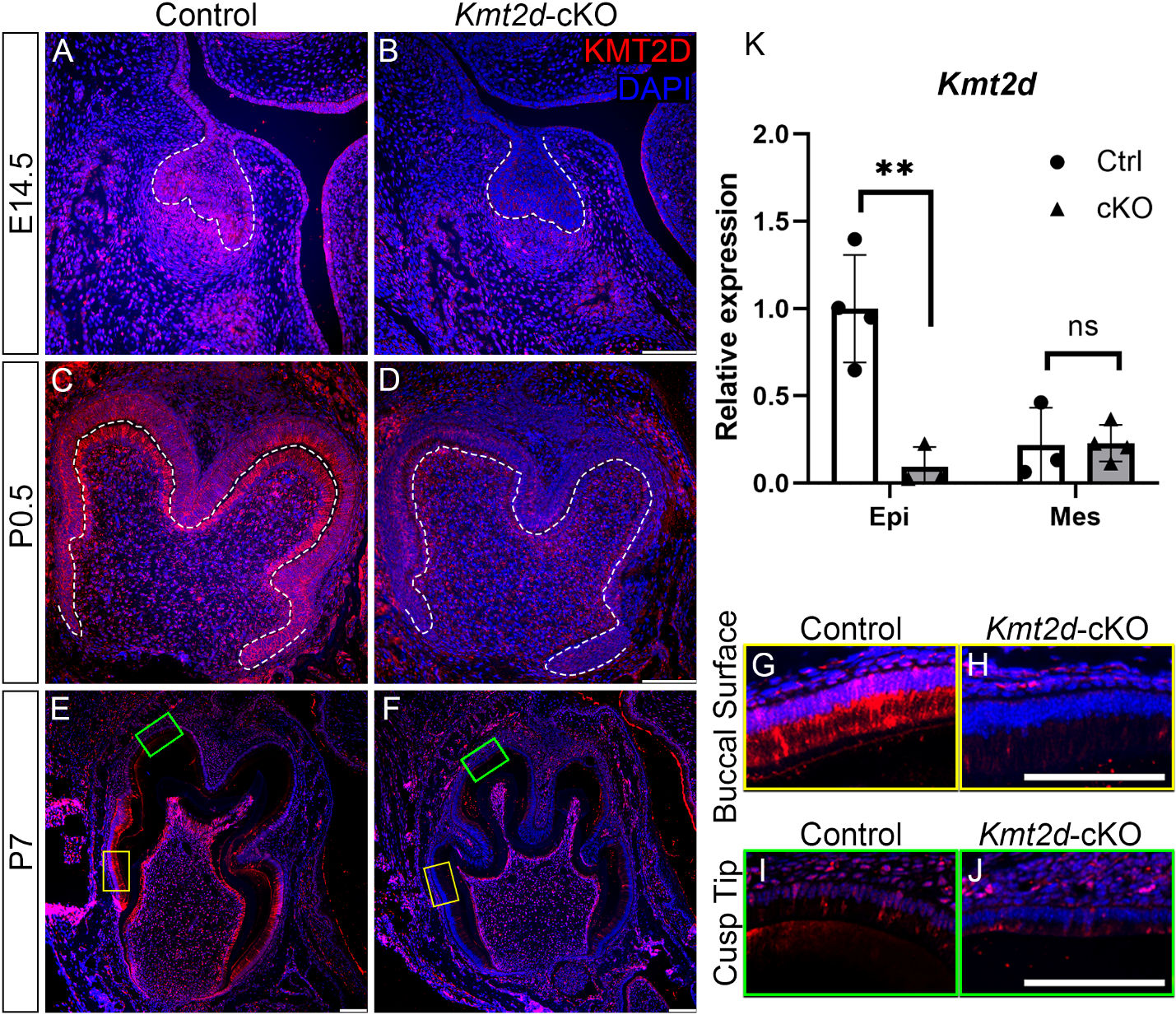

**Appendix Figure 4. *Kmt2d*-cKO tooth germs exhibit decreased KMT2D expression in the dental epithelium.** (A–J) Immunofluorescence staining for KMT2D in the first molar of control and *Kmt2d*-cKO mice at E14.5 (A, B), P0.5 (C, D), and P7 (E–J), showing markedly reduced KMT2D signals in the dental epithelium as well as dental mesenchyme of *Kmt2d*-cKO (B, D, F, H, J) compared to control (A, C, E, G, I) tooth germs. The dental epithelium-mesenchyme interface is outlined with white dashed lines (in panels A–D). Higher magnifications of the buccal surface (G, H; yellow boxes in panels E, F) and the cusp tip (I, J; green boxes in panels E, F) of the mesiobuccal ‘protoconid’ cusp both show reduced KMT2D signals in the ameloblast layer in the *Kmt2d*-cKO mice compared to controls. *n* = 4 per group. (K) qRT-PCR showing significantly decreased epithelial *Kmt2d* expression (Epi), while mesenchymal *Kmt2d* expression (Mes) remained unaffected in the *Kmt2d*-cKO first molar compared to the control. **p* < 0.05. *n* = 3–4 per group.

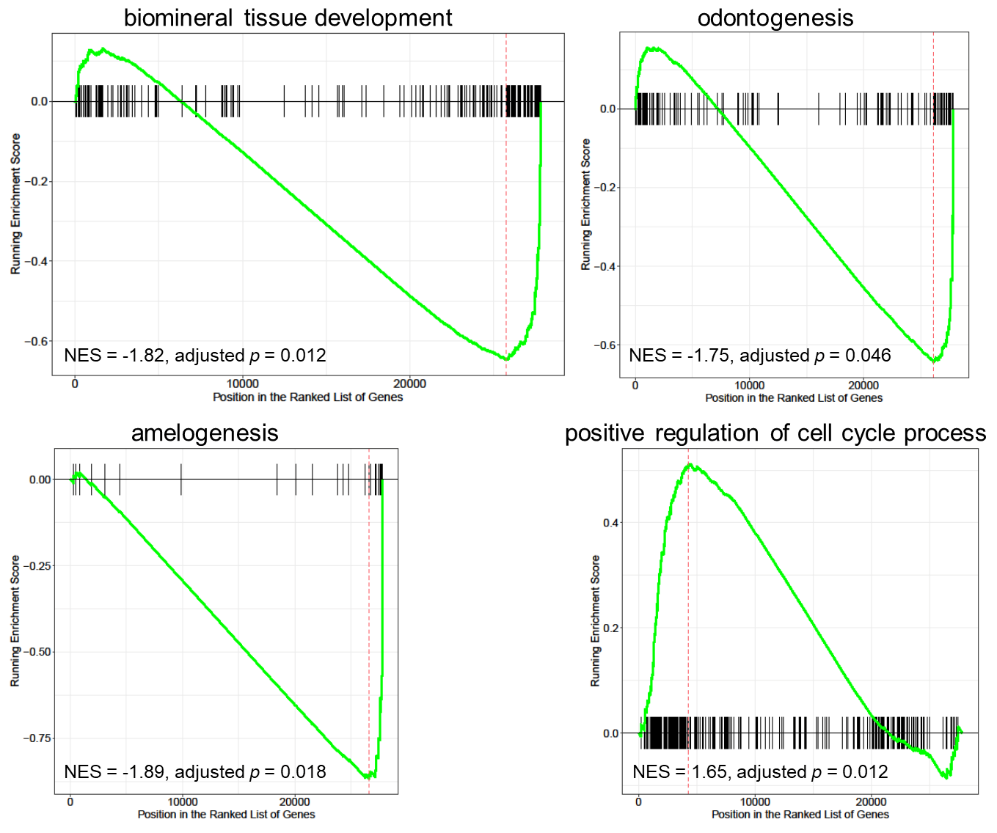

**Appendix Figure 5. Gene set enrichment (GSE) analysis-based enriched biological processes in *Kmt2d*-cKO first molars at P0.5.**

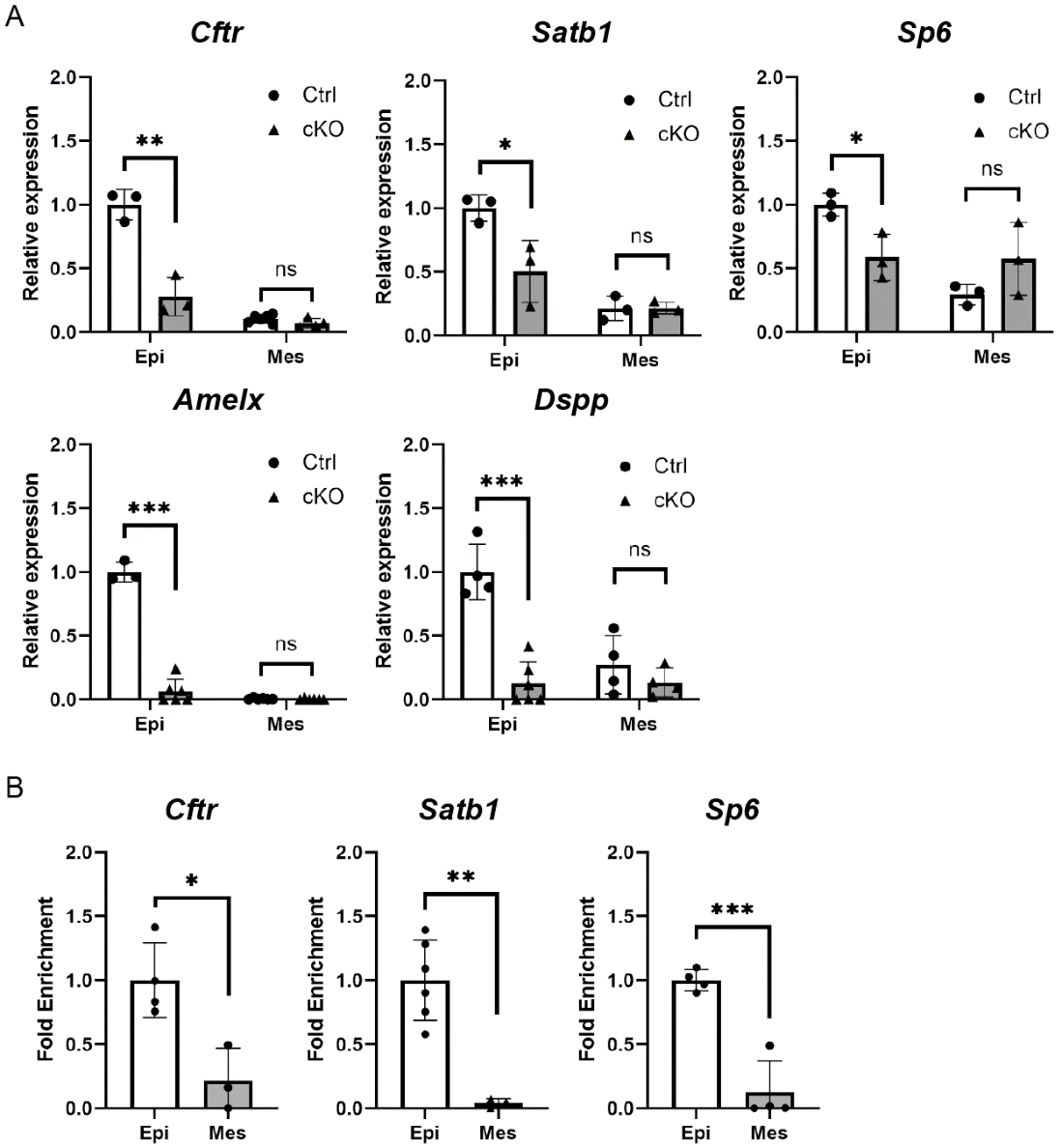

**Appendix Figure 6. Tissue-specific mRNA expression changes and fold enrichment of amelogenesis-related genes.** (A) qRT-PCR analysis of Cftr, Satb1, Sp6, Amelx, and Dspp, showing successful reduction of their mRNA expression exclusively in the Kmt2d-cKO dental epithelium, with no changes observed in the Kmt2d-cKO dental mesenchyme. (B) CUT&RUN-qPCR analysis showing specific fold enrichment of Cftr, Satb1, and Sp6 in dental epithelium compared to the dental mesenchyme. *p < 0.05; **p < 0.01; ***p < 0.001; ns, p > 0.05. n = 3–6 per group.

| *ACPT* | *COL7A1* | *GALNT3* | *MMP20* | *SLC10A7* |
| --- | --- | --- | --- | --- |
| *ADAMTS2* | *CREBBP* | *GJA1* | *MSX2* | *SLC13A5* |
| *AIRE* | *CYP27B1* | *GLB1* | *NF1* | *SLC24A4* |
| *ALDH3A2* | *DLX3* | *GNAS* | *NHS* | *SLC4A4* |
| *ALPL* | *DSPP* | *GPR68* | *OCRL1* | *SP6* |
| *AMBN* | *ENAM* | *GPR98* | *PDZD7* | *STIM1* |
| *AMELX* | *EP300* | *HCCS* | *PEX1* | *TBCE* |
| *AMTN* | *ERCC8* | *HSD17B4* | *PEX2* | *TP63* |
| *ARHGAP6* | *EVC1* | *IRX5* | *PEX26* | *TSC1* |
| *ATP6V1A* | *EVC2* | *ITGA6* | *PEX6* | *TSC2* |
| *ATR* | *FAM20A* | *ITGB4* | *PHEX* | *VDR* |
| *AVPR2* | *FAM20C* | *ITGB6* | *PITX2* | *WDR72* |
| *C4ORF26* | *FAM83H* | *KIND1* | *PLEC1* |  |
| *CELIAC1* | *FGF23* | *KL* | *PORCN* |  |
| *CFTR* | *FGF3* | *KLK4* | *PTDSS1* |  |
| *CLDN1* | *FGFR10* | *LAMA3* | *RAI1* |  |
| *CLDN16* | *FGFR2* | *LAMB3* | *RELT* |  |
| *CLDN19* | *FGFR3* | *LAMC2* | *ROGDI* |  |
| *CNNM4* | *FOXC1* | *LTBP3* | *RUNX2* |  |
| *COL17A1* | *GALNS* | *MBTPS2* | *SATB1* |  |

**Appendix Table 1. List of the 92 amelogenesis-related genes.** A total of 92 genes were compiled from previous studies (Bloch-Zupan et al. 2023; Dong et al. 2023; Smith et al. 2017; Wright et al. 2015).
